# Hijacking Cellular S-Acylation with Molecular Assemblies Rewires Oncogenic Signaling

**DOI:** 10.64898/2026.11.04.764382

**Authors:** Zhiyu Liu, Weiyi Tan, Qiuxin Zhang, Linrui Zou, Bing Xu

## Abstract

KRAS-mutant cancers are generally treated by suppressing oncogenic signaling. Here, we investigated whether endogenous lipidation machinery could instead be used to spatially perturb that signaling. The thioester-bearing peptide precursor Stb-^D^Phe-^D^Phe-SAc (**1**) potently reduced PANC-1 cell viability and formed a palmitoylated product in treated cells. Synthetic palmitoylated **1** displayed increased assembly propensity, while fluorescent analogue **2** accumulated at the Golgi in a 2-bromopalmitate-sensitive manner. Compound **1** increased plasma-membrane-associated KRAS staining and ERK/AKT phosphorylation and induced pronounced LC3B and p62 remodeling. Inhibition of palmitoylation suppressed Golgi accumulation, attenuated the LC3B response, and rescued cell viability. These findings support a model in which cellular S-acylation drives organelle-localized molecular assembly that perturbs KRAS localization, oncogenic signaling, and associated stress responses. This work identifies endogenous lipidation machinery as a synthetic handle for spatial control of cancer signaling.

---

This communication describes the use of cycling molecular assemblies (CyMA)^[1-2]^ to kill pancreatic cancer cells by altering KRAS localization and driving ERK/MAPK signaling overdose. KRAS-mutant pancreatic cancers are generally approached by suppressing oncogenic signaling. Pancreatic cancer remains one of the deadliest malignancies, and surgery, chemotherapy, and radiation are frequently insufficient for advanced disease.^[3]^ Direct KRAS inhibitors have created new therapeutic opportunities,^[4-7]^ including emerging approaches against KRAS G12X tumors,^[8]^ but adaptive and acquired resistance can limit durable responses.^[9-10]^ These limitations motivate complementary strategies that control oncogenic signaling by mechanisms other than direct inhibition.

An alternative concept is that oncogenic signaling can become a liability when it exceeds the range tolerated by a cancer cell.^[11-13]^ Disruption of SHANK3, for example, increases ERK/MAPK output and selectively impairs KRAS-mutant cells, illustrating that signaling amplification can be cytotoxic.^[14]^ Yet most approaches manipulate proteins genetically or pharmacologically; whether endogenous post-translational lipidation can be repurposed as a synthetic handle for spatially reorganizing oncogenic signaling remains largely unexplored.

Here, we asked whether CyMA^[1-2]^ could use cellular S-acylation to connect supramolecular assembly with spatial control of KRAS signaling. CyMA are a form of enzymatic self-assembly^[15-22]^ sustained by an enzymatic switch. Because the Golgi participates in KRAS localization and trafficking,^[23]^ organelle-localized assembly offered a potential route to perturb the distribution and output of the RAS-RAF-MEK-ERK network.^[24]^ Rather than assuming a predetermined death mechanism, we examined whether this chemical process would alter KRAS localization, downstream signaling, and associated cellular stress responses.

Prior CyMA studies showed reduced acquired resistance,^[1-2]^ suppression of metastasis and protumorigenic cytokine secretion in a syngeneic model, ^[1, 25]^ and high activity in tumors with elevated AKT2.^[1, 25]^

These observations suggested that dynamic intracellular assembly can influence cellular states beyond local organelle damage. PANC-1 cells provided a stringent model because they harbor KRAS^G12D^,^[26]^ exhibit aggressive and apoptosis-resistant phenotypes, and can undergo nonapoptotic death under strong stress.^[27-31]^ AKT2 is also strongly expressed in PANC-1 and contributes to metastatic progression.^[32]^

We therefore designed thioester-bearing CyMA precursors to test a prediction gap between two fields: S-acylation is typically viewed as a mechanism that controls endogenous protein localization, whereas supramolecular assembly is usually used to create local intracellular structures. We hypothesized that coupling these processes could transform endogenous lipidation machinery into a synthetic spatial actuator Our results support this possibility. Compound **1** potently reduced PANC-1 cell viability and yielded a palmitoylated product in treated cells; the synthetic palmitoylated product displayed increased assembly propensity. Fluorescent analogue **2** accumulated at the Golgi in a 2-bromopalmitate (2-BP)-sensitive manner, while compound **1** increased plasma-membrane-associated KRAS staining, ERK/AKT phosphorylation, and pronounced LC3B and p62^[33]^ remodeling. Inhibiting palmitoylation reduced Golgi accumulation, attenuated the LC3B response, and rescued viability. Together, these observations support a model in which cellular S-acylation drives organelle-localized molecular assembly that perturbs KRAS localization, oncogenic signaling, and associated stress responses. This framework treats endogenous lipidation machinery not only as a biological pathway to inhibit, but also as a synthetic handle for spatial control of cancer signaling.

**Scheme 1.**
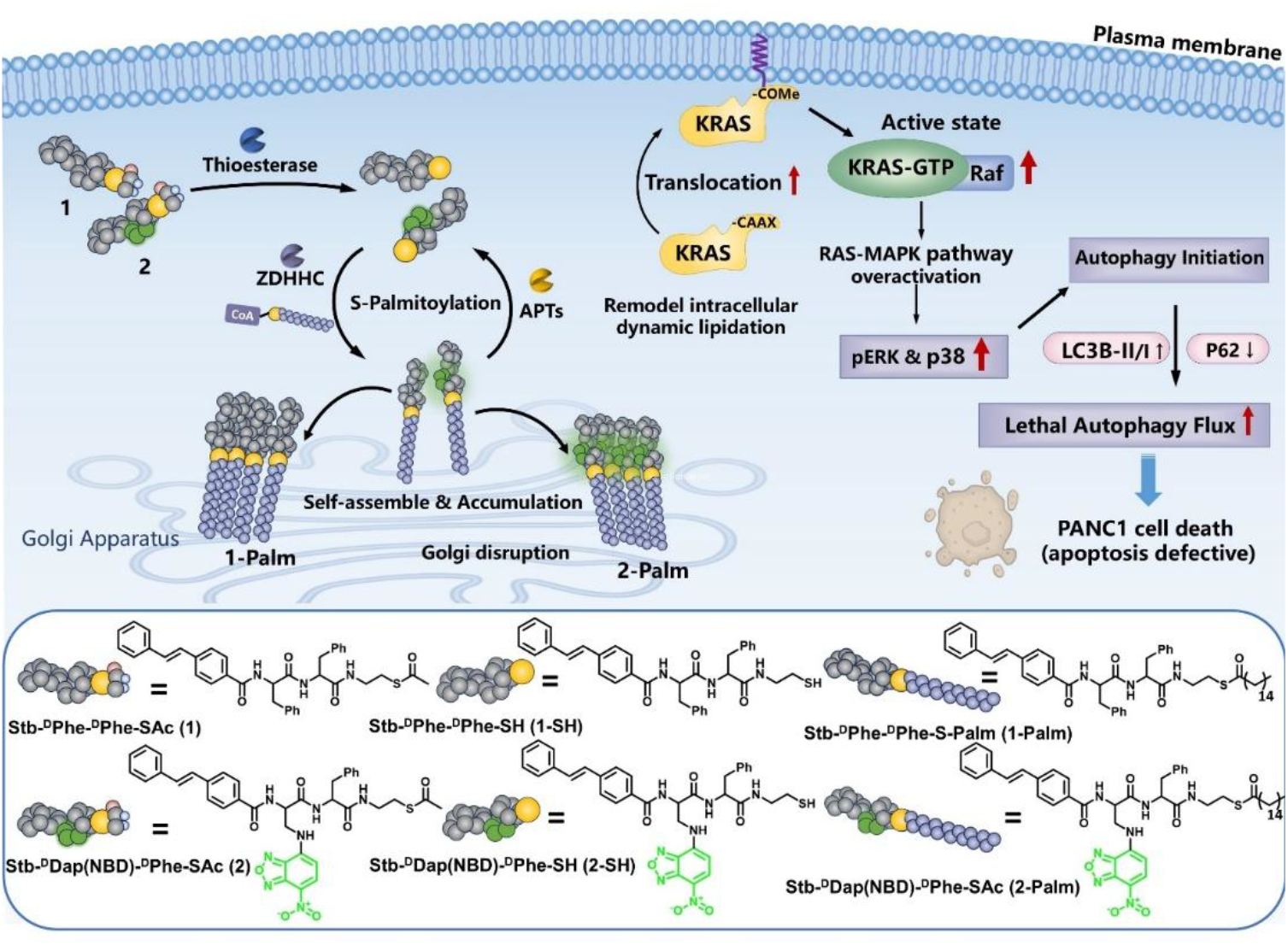
Proposed mechanism by which cycling molecular assemblies (CyMA) induce pancreatic cancer cell death. An enzyme-regulated cycle of thioesterase-mediated deacylation and ZDHHC-mediated S-palmitoylation promotes CyMA accumulation and assembly at the Golgi. The resulting Golgi perturbation redistributes KRAS to the plasma membrane, enhances KRAS–RAF–MEK–ERK signaling, and drives excessive autophagic flux, leading to PANC-1 cell death.

We designed two peptide-based CyMA precursors (Scheme 1), each consisting of three key modules: (i) an N-terminal stilbene (Stb) capping group, (ii) a diphenylalanine backbone^[34]^, and (iii) a C-terminal thioester. The thioester enables enzyme-regulated, reversible S-palmitoylation through the opposing activities of thioesterases^[35]^ and palmitoyltransferases (ZDHHCs).^[36]^ Precursor **1** contains D-diphenylalanine to promote self-assembly, while precursor **2** incorporates an NBD fluorophore to enable intracellular imaging.^[37]^

After synthesizing **1** and **2** (Schemes S2 and S3), we first evaluated their antiproliferative potency in PANC-1 cells. Both molecules significantly inhibited PANC-1 cell growth (Figures S1 and S2). **1** showed potent activity, with a GI_50_ of approximately 130 nM and a GI_90_ around 700 nM at 24 h (Figure 1A), whereas **2** exhibited a GI_50_ and of 4 µM and a GI_90_ of 25 µM. This higher potency of **1** likely reflects its stronger self-assembly propensity, conferred by the combined Stb and D-diphenylalanine motifs. At 100 nM, **1** produced no detectable cytotoxicity in HepG2 hepatocyte-like cells (Figure S3), indicating selectivity within this concentration range. We next compared CyMA with known KRAS inhibitors MRTX1133,^[38]^ sotorasib, and daraxonasib in PANC-1 cells.^[39]^ CyMA showed two to three orders of magnitude greater antiproliferative potency than these KRAS inhibitors under the tested conditions (Figures S4, S5, S15, and S16).

**Figure 1.**
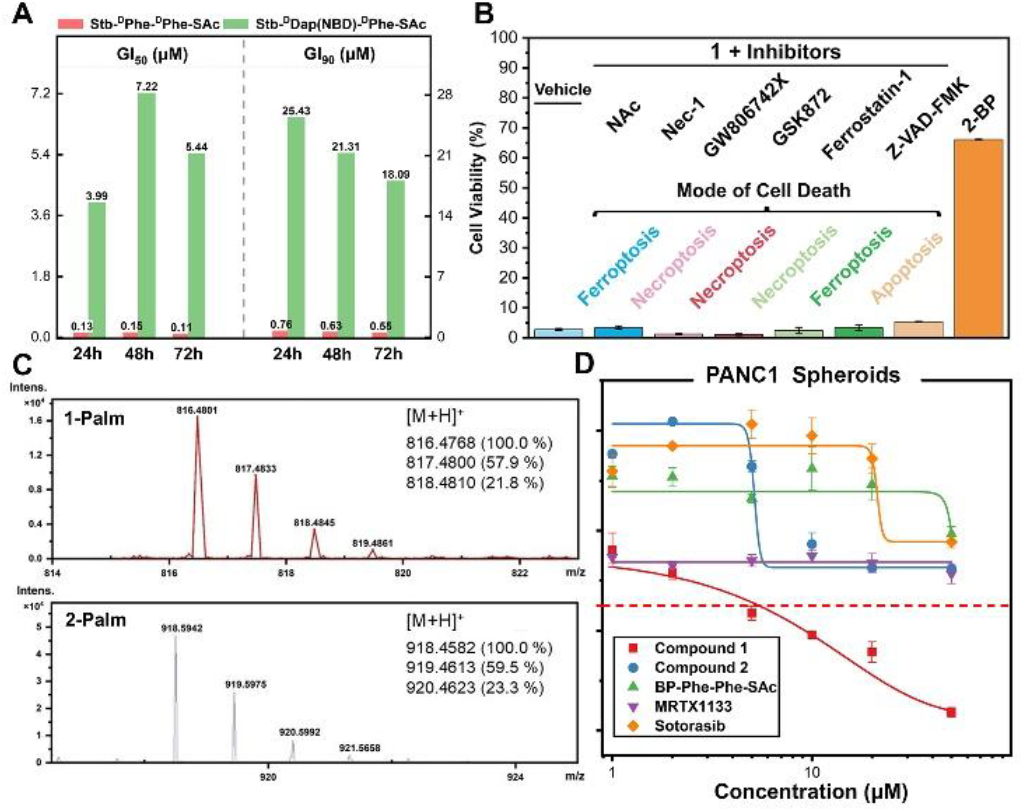
(A) GI_50_ and GI_90_ values of **1** and **2** in PANC-1 cells after 24, 48, and 72 h of treatment. (B) Viability of PANC-1 cells treated with **1** (1 μM) for 24 h in the presence or absence of inhibitors of canonical cell-death pathways: N-acetylcysteine (NAC, 100 μM), necrostatin-1 (Nec-1, 100 μM), GW806742X (10 μM), GSK872 (10 μM), ferrostatin-1 (10 μM), Z-VAD-FMK (100 μM), or 2-bromopalmitate (2-BP, 10 μM). DMSO served as the vehicle control; n = 3. (C) High-resolution LC– MS spectra of palmitoylated **1** (**1**-Palm) and palmitoylated **2** (**2**-Palm) extracted from lysates of PANC-1 cells treated with 1 or 2, respectively, for 2 h. (D) Viability of PANC-1 spheroids after treatment with **1, 2**, BP-^D^Phe-^D^Phe-SAc, MRTX1133, or sotorasib for 72 h; n = 3.

To define the mechanism underlying CyMA-induced cytotoxicity, we co-treated PANC-1 cells with **1** and a panel of pharmacological inhibitors, including inhibitors of canonical cell death pathways.^[40-41]^ Among the compounds tested, only 2-bromopalmitate (2-BP) substantially rescued cell viability, restoring it to over 65% (Figures 1B and S6). This result implicates S-palmitoylation in the cytotoxic activity of **1**. We further tested this requirement using analogues **3** and **4**, in which replacement of the thioester with a carboxyl ester prevented S-palmitoylation and markedly reduced cytotoxicity (Figure S7). Together, these results establish reversible S-palmitoylation as a critical determinant of CyMA formation and biological activity.

To probe this mechanism further, we analyzed lysates from PANC-1 cells treated with **1** or **2** for 2 h using high-resolution liquid chromatography–mass spectrometry (HRMS). Theanalysis detected new peaks corresponding to palmitoylated **1** (**1**-Palm) or palmitoylated **2** (**2**-Palm), with isotope distributions consistent with the assigned products (Figures 1C & S8). We next measured the critical micelle concentrations (CMC) of **1, 2, 5** (deacylated **1)**, and **6** (palmitoylated **1**) (Figure S9-S12). Compound **6** displayed the lowest CMC and formed aggregates at low concentration, as confirmed by transmission electron microscopy (TEM).

(Figure S13). These results demonstrate that palmitoylation markedly increases the assembly propensity of **1**.

To define the structural requirements for activity, we systematically varied the linkage between the two aromatic rings of **1**, synthesized a series of analogues, and evaluated their effects on PANC-1 cell growth. The analogs exhibited distinct GI_50_ values (Figure S14), demonstrating the sensitivity of biological activity to subtle structural alterations. These linkages modifications likely alter molecular rigidity and polarity, and membrane interactions, thereby affecting CyMA formation at the Golgi and the resulting cytotoxic potency.

We next evaluated the activity of CyMA precursors and clinically used drugs in 3D PANC-1 spheroids (Figure 1D). Among the compounds tested, only **1** showed strong cytotoxicity, reducing spheroid viability to below 10%. None of the other compounds reduced viability below 50% at the concentration up to 200 µM. These results agree with the cytotoxicity profiles observed in 2D cell cultures and demonstrate the potent activity of **1** in a physiologically more relevant tumor model.

We used fluorescent analogue **2** to visualize the intracellular localization of CyMA. Confocal fluorescence microscopy showed that **2** accumulated predominantly at the Golgi and strongly colocalization with a Golgi marker,^[42]^ with a mean correlation coefficient of 0.89 (Figure 2A). We then treated PANC-1 cells with **2** alone or in combination with the palmitoylation inhibitor 2-BP for 30 minutes. In cells treated with **2** alone, fluorescence increased over time and concentrated at the Golgi. By contrast, 2-BP abolished this accumulation (Figures 2B and S17). Line-scan analysis at 30 min revealed a pronounced fluorescence at the Golgi region, further confirming Golgi localization. Golgi-associated fluorescence increased within 20 min and remained significantly higher than that observed after cotreatment with 2-BP (Figure 2C,D). In the absence of 2-BP, **2** remained at the Golgi for an extended period (Figure 2E). Together, these results indicate that reversible S-acylation promotes the accumulation and prolonged retention of CyMA at the Golgi.

**Figure 2.**
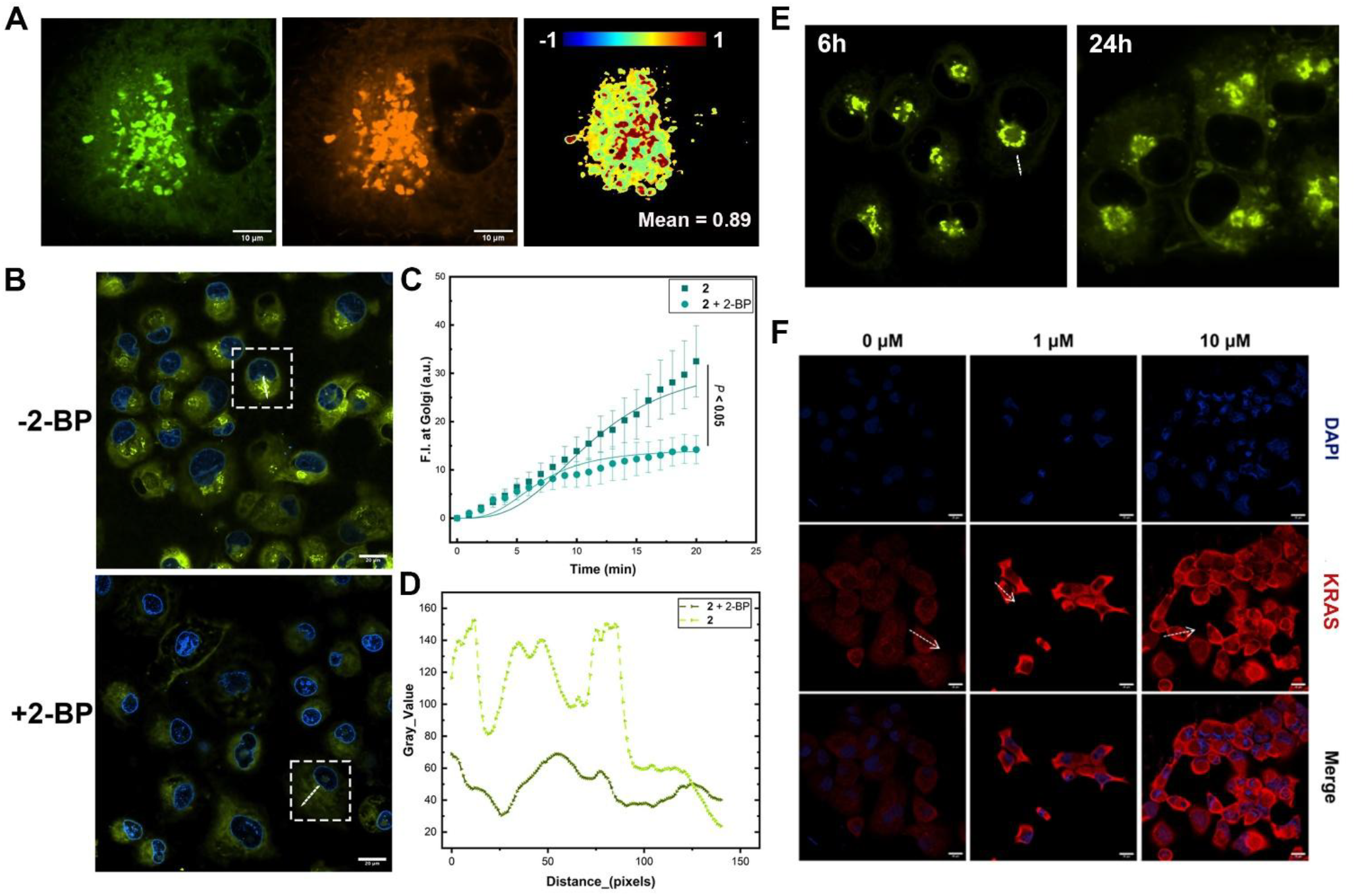
(A) Confocal laser scanning microscopy (CLSM) images of PANC-1 cells treated with **2** (10 μM) and a Golgi tracker^[42]^ for 2 h. The colocalization map shows a mean Pearson correlation coefficient of 0.89. Scale bars, 20 μm. (B) CLSM images of PANC-1 cells treated with **2** (2 μM) for 30 min in the absence or presence of 2-bromopalmitate (2-BP, 20 μM). Nuclei were stained with Hoechst 33342. Dashed boxes indicate regions used for line-scan analysis. Scale bars, 20 μm. (C) Time-dependent Golgi-associated fluorescence intensity in PANC-1 cells treated with **2** alone or with **2** and 2-BP; n = 6. (D) Line-scan fluorescence profiles across the Golgi regions indicated in panel B. (E) CLSM images showing the intracellular retention of **2** (2 μM) in PANC-1 cells after 6 and 24 h. (F) Immunofluorescence images showing KRAS localization in PANC-1 cells after treatment with **1** at the indicated concentrations for 12 h. KRAS was detected with an Alexa Fluor 647-conjugated secondary antibody, and nuclei were stained with Hoechst 33342. Scale bar = 20 μm.

The carboxyl ester analog of **2**, precursor **4** (Stb-^D^Dap(NBD)-^D^Phe-OAc), showed minimal Golgi localization, with a mean correlation coefficient of 0.12 (Figures S18 and S19). This loss of localization further supports a requirement for thioesterase- and ZDHHC-mediated reversible S-palmitoylation in Golgi targeting. Notably, treatment with **1** also increased KRAS accumulation at the plasma membrane in a concentration-dependent manner (Figures 2F & S20), while cotreatment with 2-BP eliminated this phenomenon (Figure S21), suggesting that CyMA shifts the lipidation-dependent trafficking equilibrium of KRAS toward plasma-membrane retention.

We next investigated how CyMA induces PANC-1 cell death by examining signaling and trafficking protein after treatment with **1**. Rab11A and Rab11B levels remain unchanged across the tested concentrations (Figure 3A), indicating that **1** does not broadly suppress Rab11-dependent recycling or secretory trafficking.^[43]^ In contrast, **1** increased the levels of phosphorylated ERK (pERK) and phosphorylated AKT (pAKT), as well as the ratios of pERK/ERK and pAKT/AKT (Figure S22). These finding are consistent with increased KRAS localization at PM, where KRAS activates downstream ERK and AKT signaling.^[39]^

**Figure 3.**
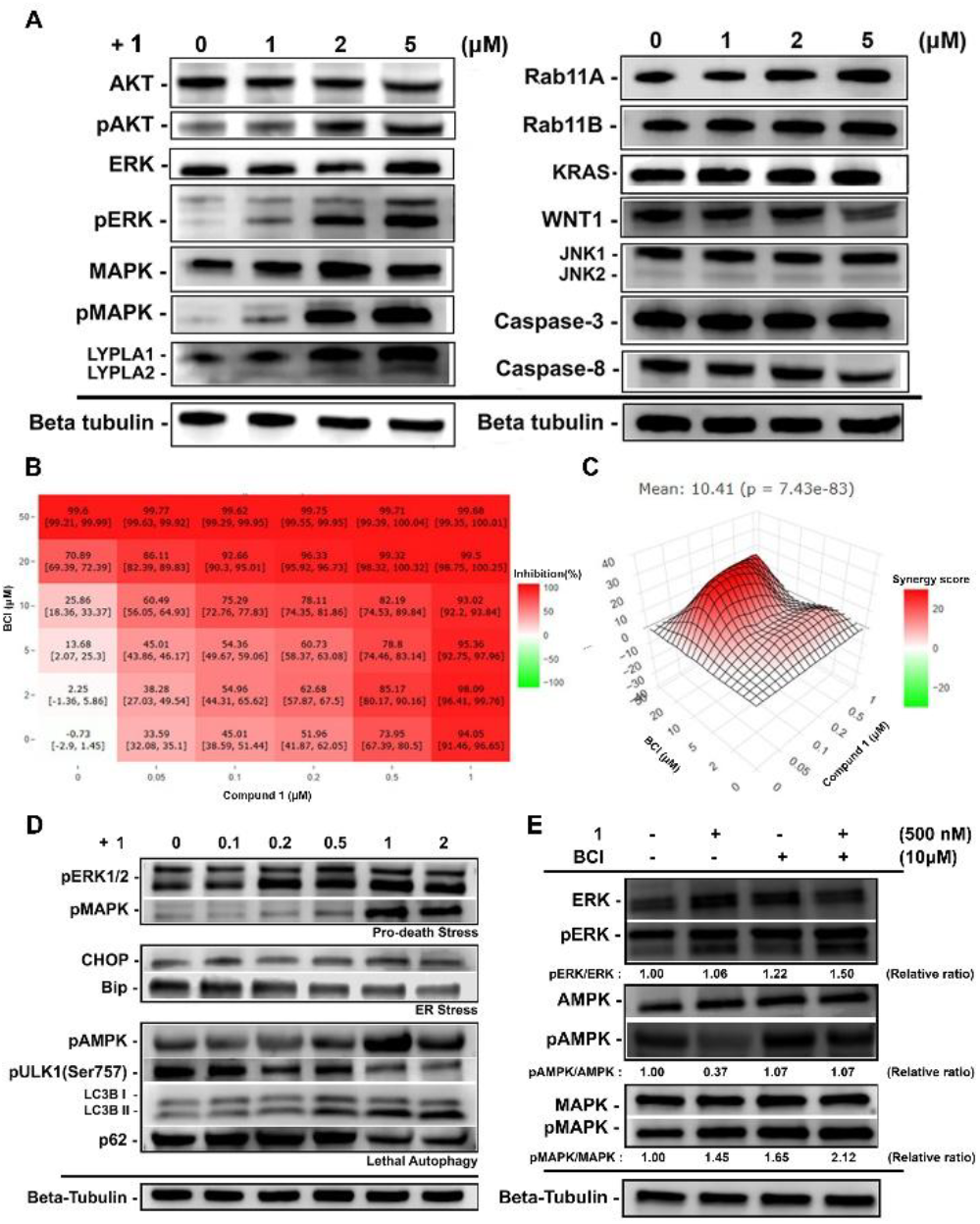
(A) Immunoblot analysis of proteins associated with growth and stress signaling, reversible S-palmitoylation, and intracellular trafficking in PANC-1 cells treated with **1** at the indicated concentrations for 12 h. (B) Combination dose–response matrix showing inhibition of PANC-1 cell viability after cotreatment with **1** and the DUSP1/6 inhibitor BCI. (C) Highest-single-agent (HSA) synergy analysis of the combination in panel B. Positive synergy occurred across a broad concentration range and reached a maximum at intermediate concentrations of both compounds. (D) Immunoblot analysis of proteins associated with ER stress and autophagy in PANC-1 cells treated with **1** at the indicated concentrations for 12 h. (E) Immunoblot analysis of ERK/MAPK and AKT signaling in PANC-1 cells treated with **1** (500 nM), BCI (10 μM), or their combination for 24h.

Immunocytochemistry and western blotting further showed that **1** increased AMPK phosphorylation while decreasing ULK1 phosphorylation at Ser757, indicating activation of the AMPK–ULK1 axis and relief of mTORC1-mediated inhibition (Figure 3D). **1** also increased LC3B levels in a concentration-dependent manner (Figures 3D and S23) and markedly reduced p62 abundance, consistent with increased autophagic flux. Together, these results indicate that CyMA-driven retention of KRAS at PM produces sustained oncogenic signaling and cellular stress, which activate the AMPK– ULK1 pathway and induce autophagy.

In PANC-1, sustained co-activation of p38 and ERK shifts cell stress towards autophagy-dependent death. Chloroquine (CQ), which inhibits lysosomal degradation and blocks completion of autophagic flux, increased the cytotoxicity of **1** and **2**. In contrast, ULK1inhibition partially rescued PANC-1 cells (Figures S24 and S25). Together, these findings indicate that CyMA-induced cytotoxicity requires autophagy and highlight the context-dependent role of autophagy: it initially supports stress adaptation but becomes death-promoting under persistent CyMA-induced signaling stress. Treatment of 2-BP reverses this phenotype by inhibiting the S-palmitoylation of **1**, attenuating ERK activation, and restoring autophagic homeostasis^[44]^ (Figure S26). These results show that **1** kills PANC-1 cells by inducing palmitoylation-dependent, sustained ERK signaling, whereas inhibition of palmitoylation interrupts this signaling program and preserves cell viability.

Treatment with **1** hardly altered the total abundance of Rab11A/B, KRAS, WNT1, JNK1/2 (Figure 3A), and did not induce cleavage of caspase-3/8 (Figure S27), indicating that cytotoxicity arises primarily from changes in protein localization and posttranslational signaling rather than altered protein expression. This interpretation is consistent with sustained ERK and p38 phosphorylation, relief of inhibitory control over ULK1, and increased autophagy flux in PANC-1 cells.

To examine the contribution of MAPK negative-feedback regulation, we cotreated PANC-1 cells with **1** and BCI, an inhibitor of the DUSP1 and DUSP6. The combination produced strong overall efficacy (CSS = 93.6) and synergistic interactions across multiple reference models, including an HSA synergy score of 10.41 (Figures 3B,C and S28). The strongest synergy occurred at sub-saturating concentrations of both compounds. Mechanistically, BCI further enhanced ERK/MAPK activation induced by **1** (Figure 3E), indicating that inhibition of DUSP-mediated adaptive feedback amplifies CyMA-induced MAPK signaling stress. Conversely, pharmacological inhibition of MEK by trametinib partially attenuated the antiproliferative activity of **1** (Figure S29), further supporting the involvement of ERK/MAPK signaling in the cellular response elicited by CyMA. Moreover, CyMA maintained its activity after the repeated-treatment protocol (Figure S30), suggesting a low propensity to induce drug resistance. By perturbing reversible lipidation and KRAS signaling dynamics, CyMA may provide a strategy to overcome adaptive resistance.

In summary, this study identifies cellular S-acylation as a synthetic handle for directing dynamic molecular assembly within cancer cells. By harnessing the cell-intrinsic reversible S-palmitoylation machinery,^[45-46]^ CyMA promotes intracellular assembly at specific subcellular sites, perturbs protein localization and signaling, and potently suppresses pancreatic cancer cell viability through an autophagy-dependent mechanism.^[47]^ The accompanying increase in LYPLA1 expression (Figure 3A) further suggests that CyMA disturbs the regulated balance between S-palmitoylation and depalmitoylation, thereby altering lipidation-dependent trafficking and cellular homeostasis.

These findings establish dynamic peptide assembly as a strategy for perturbing reversible protein lipidation in cancer cells.^[48]^ More broadly, they define a molecular design principle in which organelle-localized supramolecular assemblies reprogram oncogenic signaling, cellular stress responses, and autophagy to alter cell fate. Such an approach may provide a complementary strategy for converting cancer-associated lipidation and signaling pathways into selective vulnerabilities in treatment-refractory pancreatic cancer.

## Supporting Information

This material is available free of charge via the Internet at detailed experimental procedures and materials; HPLC and MS spectra for individual compounds; CLSM and Immunocytochemistry images, cell viabilities, and chemical structures of the compounds (PDF). The authors have cited additional references within the Supporting Information.

## Acknowledgements

We thank the Brandeis Light Microscopy Facility (RRID:SCR_025892) for assistance. This work is partially supported by NIH (CA142746).

## Entry for the Table of Contents

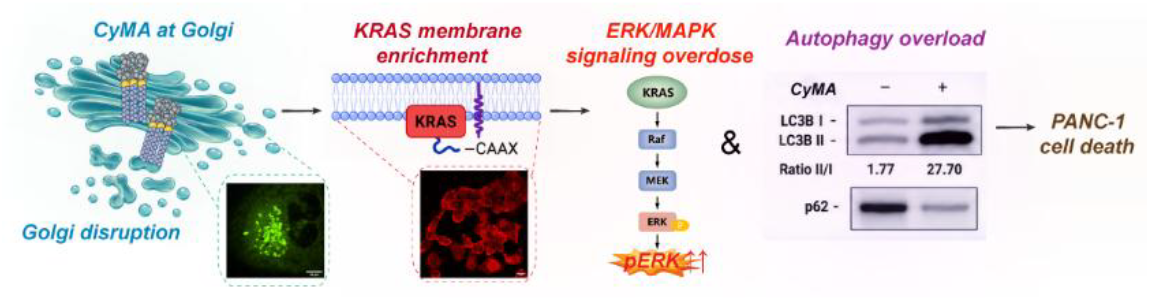

Cycling molecular assemblies (CyMAs) harness cellular reversible S-palmitoylation to accumulate at the Golgi and redirect KRAS localization. Instead of inhibiting oncogenic signaling, CyMA amplify ERK/MAPK activity beyond the cellular tolerance threshold, causing excessive autophagic flux and potent death of pancreatic cancer cells.

