## Supplemental information for "Hijacking Cellular S-Acylation with Molecular Assemblies Rewires Oncogenic Signaling"

**Table of Contents**

Supplemental Experimental Procedures .......... 3

Supporting figures .......... 15

### Supplemental Experimental Procedures

#### Materials

2-Cl-trityl chloride resin (1.0 mmol/g), Fmoc-D-Phe-OH, Fmoc-D-Ala-OH, and HBTU were obtained from GL Biochem (Shanghai, China). N, N-diisopropylethylamine (DIPEA), 2-Mercaptoethanol, chloroquine diphosphate salt, and solvents were obtained from Fisher Scientific. 4-Chloro-7-nitrobenzofurazan was obtained from Alfa Aesar. Fmoc-$^{D}$Dap(Boc)-OH, 2-Aminoethyl acetate hydrochloride, 4-(phenylethynyl)benzoic acid and cysteamine hydrochloride were purchased from AmBeed. 1-[(2-Aminoethyl)sulfanyl]ethan-1-one hydrobromide, 2-bromohexadecanoic acid, 4-(Phenylazo)benzoic acid were purchased from Sigma-Aldrich. Acetyl chloride and palmitoyl chloride were purchased from TCI America. Biphenyl-3-carboxylic acid was purchased from 1PlusChem. 4-Benzylbenzoic acid, 4-(2-phenylethynyl) benzoic acid were purchased from Enamine. (E)-4-styrylbenzoic acid was purchased from AChemBlock. Necrostatin-1, acetylcysteine, GW806742X, GSK872, Ferrostain-1, Z-VAD-FMK, MRTX1133, Sotorasib, BCI, Daraxonrasib and SBI-0206965 were purchased from MedChemExpress. All the chemical reagents and solvents were used as received from commercial sources without further purification.

Goat anti-rabbit IgG (H+L) secondary antibody was purchased from Invitrogen. Alexa Fluor™ 647 conjugate (Catalog # C34778) was purchased from Invitrogen. TWEEN® 20, and Immobilon™-FL PVDF membranes (catalog # IPFL00005) were purchased from Millipore Sigma. LYPLA1 polyclonal antibody (Catalog # PA5-101405), LYPLA2 polyclonal antibody (Catalog # PA5-27653), mTOR polyclonal antibody (Catalog # PA5-34663), Phospho-ERK1/ERK2 polyclonal antibody (Catalog # 44-680G), ERK1/ERK2 monoclonal antibody (Catalog # MA5-15134), Phosphop38 MAPK monoclonal antibody (Catalog # MA5-15182), p38 MAPK recombinant rabbit monoclonal antibody (Catalog # MA5-41213), HRP (catalog # 31460), goat anti-rabbit IgG (Heavy chain), superclonal™ recombinant secondary antibody, Alexa Fluor™ 647 (catalog # A27040), SuperSignal™west pico PLUS chemiluminescent substrate (catalog # 34579), SuperBlock™ (TBS) blocking buffer (catalog # 37535), Restore™ PLUS western blot stripping buffer (catalog # 46428), and Halt™ protease inhibitor cocktail (100X) (catalog # 78430) were purchased from Invitrogen. KRAS polyclonal antibody (Catalog # 12063-1-AP), JNK polyclonal antibody (Catalog # 10023-1-AP), Goat anti-rabbit IgG H&L (Alexa Fluor® 647) (Catalog # ab150079), rabbit recombinant monoclonal LC3B antibody (Catalog # ab192890), anti-pan-AKT antibody (Catalog # ab8805), anti-pAKT antibody (Catalog # ab192623), anti-beta Actin antibody (Catalog # ab8227) were purchased from abcam. CHOP antibody (Catalog # 5554), Bip antibody (Catalog # 3183), Caspase-3 antibody (Catalog # 9662), Caspase-8 antibody (Catalog # 5554), K-Ras antibody (Catalog # 33197), Phospho-ULK1 (Ser757) antibody (Catalog #6888), SQSTM1/p62 antibody (Catalog #5114) and Beta-Tubulin antibody (Catalog # PA5-31348) were purchased from Cell Signaling. Rab11A polyclonal antibody (Catalog # 0229-1-AP), Rab11B polyclonal antibody (Catalog # PA5-31348), WNT1 polyclonal antibody (Catalog # PA5-34776) were purchased from ThermoFisher.

Dulbecco's Modified Eagle Medium (DMEM), fetal bovine serum (FBS), Opti-MEM, $CO_2$ Independent Medium and penicillinstreptomycin (PS) were purchased from Gibco.Nunclon™ Sphera™ 96-Well, Nunclon Sphera-Treated, U-Shaped-Bottom Microplate (Catalog # 174929) were purchased from Thermo Scientific. Inc. CellTiter-Glo® 3D Cell Viability Assay S9 (Catalog # G9681) was purchased from Promega.

### Instruments

All precursors and compounds were purified by a reverse phase HPLC (Agilent 1100 Series) equipped with an XTerra C18 RP column. HPLC grade acetonitrile (0.1% TFA) and HPLC grade water (0.1% TFA) were used as the eluents. LC-MS spectra were obtained with a Waters Acquity Ultra Performance LC with Waters MICROMASS detector, and a Bruker Elute PLUS UHPLC with a Bruker tims TOF Pro. Fluorescence images were captured using a Nikon AX-R Resonant Confocal System. TEM images were conducted on Morgagni 268 transmission electron microscope.

### Methods

#### Synthesis of Fmoc-$^{D}$Dap(NBD)-OH

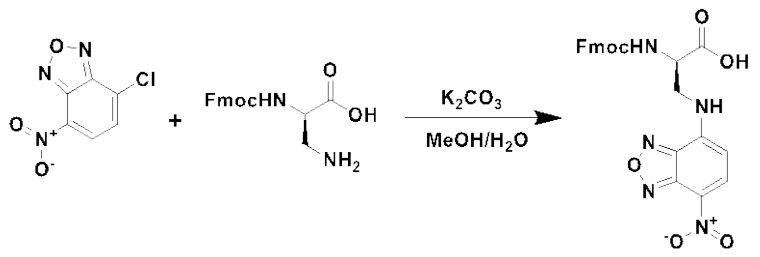


**Scheme S1.** Synthetic procedure of Fmoc-$^{D}$Dap(NBD)-OH.

Fmoc-$^{D}$Dap-OH (1 mmol, 326 mg) and potassium carbonate (3 mmol, 414 mg) were mixed in a 10 mL aqueous solution, NBD-Cl (1.2 mmol, 240 mg) in 15 mL MeOH was added dropwise with stirring under nitrogen gas protection. After stirring at room temperature for 6 hours, the methanol was removed using a rotary evaporator and the residual solution was acidified to pH 3 using 1 M HCl. The acidic aqueous solution was then extracted by diethyl ether. The combined organic solution was dried over anhydrous sodium sulfate and concentrated by rotary evaporator. The resulting dark-yellow powder (Fmoc-$^{D}$Dap(NBD)-OH) was used directly for solid phase peptide synthesis.

**Synthesis of CyMA 1, 3 and derivatives**

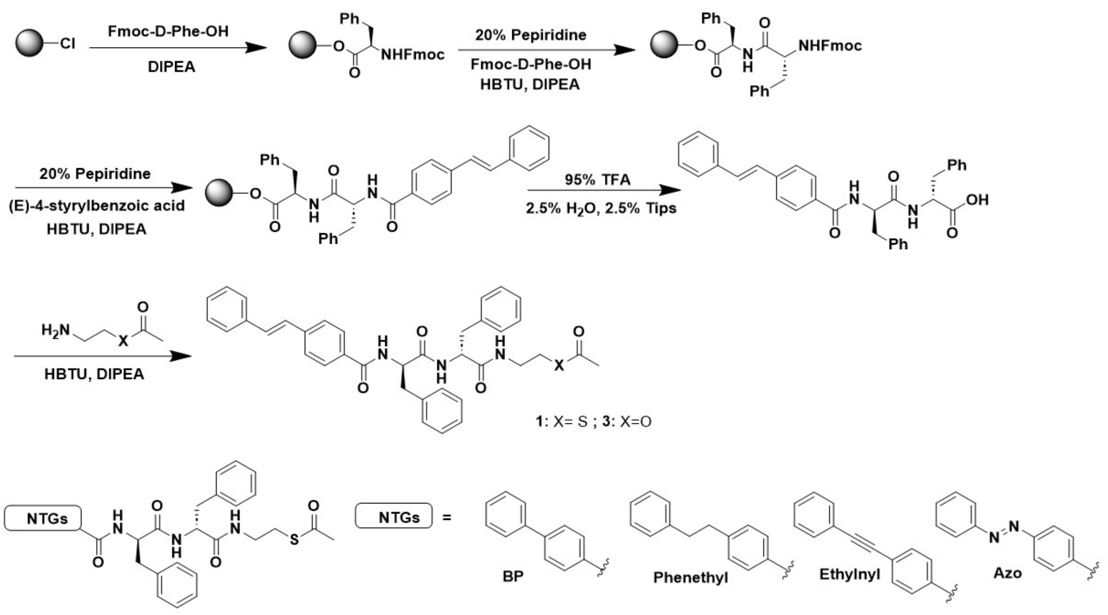


**Scheme S2.** Synthetic procedure of compound **1** and **3 and derivatives.**

We used standard Fmoc chemistry for solid phase peptide synthesis (SPPS) with 2-chlorotrityl chloride resin and Fmoc-protected amino acids with appropriately protected side chains. Briefly, the 2-Cl resin (0.5 g) was swelling in dry DCM for 1 hour. Next, 2 equivalents of Fmoc-$^{D}$Phe-OH (1 mmol, 389 mg) and DIPEA in DCM were added into the SPPS reactor and incubated with the resin for 2 hours at room temperature. After discarding the liquid phase, the resin was washed three times with DCM. Subsequently, blocking buffer (DCM: MeOH: DIPEA = 17:2:1) was added into resin for 15 mins and then washed three times with DMF. The Fmoc group was removed with 20% piperidine in DMF, and the next Fmoc-protected amino acid (Fmoc-$^{D}$Phe-OH, 1 mmol) was coupled to the free amino group using

HBTU (1 mmol) as the coupling reagent. The N-terminus of the peptide was capped with (E)-4-styrylbenzoic acid on the resin. The peptide chain was cleaved from the resin by 95% TFA (95% TFA, 2.5% TIPS, 2.5% $H_2O$) for 1 hour. After solvent removal via rotary evaporation, 30 mL of dry diethyl ether was added to the residual solution, followed by centrifugation at 5000 rpm for 8 minutes. The resulting solid products were weighed out 0.1 mmol, followed by adding 0.12 mmol HBTU and 0.12 mmol 1-[(2-Aminoethyl)sulfanyl]ethan-1-one hydrobromide for CyMA **1** or 2-Aminoethyl acetate hydrochloride for CyMA **3**, all the reagents were dissolved in 2 mL DMF with pH around 8-9 adjusted by DIPEA, reacted overnight. After solvent removal via air dry, the resulting solid products were further purified using RP-HPLC (Scheme S2). Four different NTGs including 2-bromohexadecanoic acid, 4-(Phenylazo)benzoic acid, 4-Benzylbenzoic acid, 4-(2-phenylethynyl)benzoic acid were used for replacement of 4-styrylbenzoic acid to get different derivatives.

**Synthesis of 2, 4**

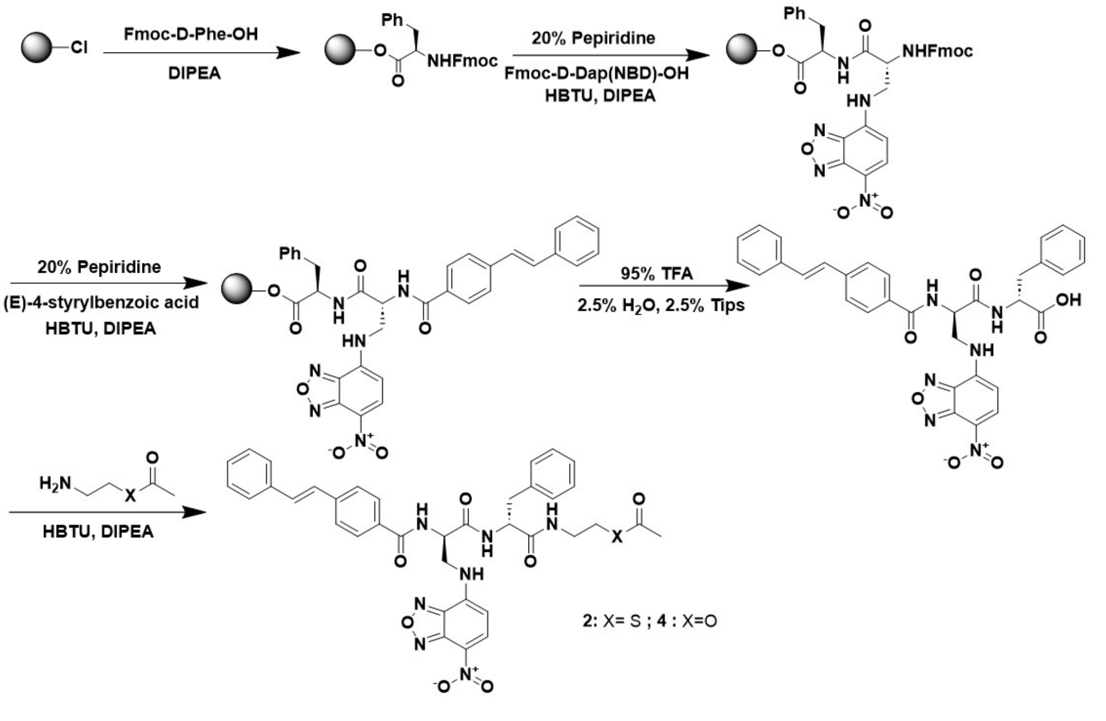

**Scheme S3.** Synthetic procedure of the compound **2**, **4** and derivatives.

The synthetic procedure of CyMA-i is almost the same as CyMA-d, only difference is that the second amino acid Fmoc-$^{D}$Phe-OH was replaced by Fmoc-$^{D}$Dap(NBD)-OH. Also, 1-[(2-Aminoethyl)sulfanyl]ethan-1-one hydrobromide was used for CyMA **2** with thiol-acyl at C-terminal and 2-Aminoethyl acetate hydrochloride was used for CyMA **4** with O-ester at C-terminal (Scheme S3).

**Synthesis of 1-**SH (**5**)**, 1-**Palm (**6**)

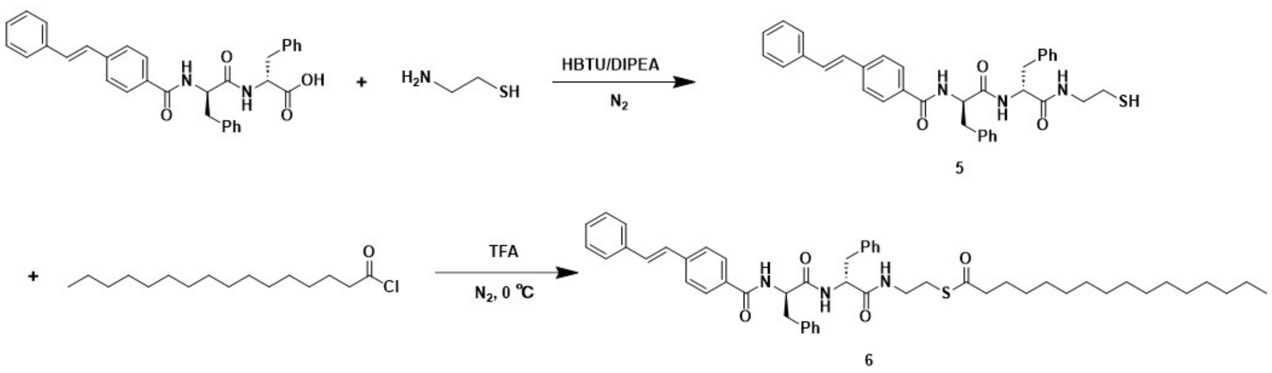


**Scheme S4.** Synthetic procedure of compounds **5** and **6**.

The Stb-$^{D}$Phe-$^{D}$Phe-OH (0.1 mmol, 51.8 mg) from procedure in scheme S2 and HBTU (0.12 mmol) were dissolved in 2 mL DMF and stir for 30 mins for activation. Then cysteamine hydrochloride (0.2 mmol) was added under protection of $N_2$, followed by adding enough DIPEA to adjust the pH to 8, react overnight (Scheme S4). After the reaction finished, the solvent was dried by $N_2$, the resulting solid products were further purified with RP-HPLC to get **1**-SH (**5**). Then 1-SH (0.01 mmol, 5.8 mg) was dissolved in 2 mL TFA and stir in ice bath and 10 equivalents of palmitoyl chloride was added dropwise into the solution under the protection of $N_2$, react overnight. After solvent removal via air dry, the resulting solid products were further purified with RP-HPLC to get **1-**Palm (**6**) (Scheme S4).

**Cell culture**

PANC-1 cells were purchased from ATCC. Human pancreatic ductal epithelial cells (HPDEC) were purchased from AddexBio. PANC-1 cells were cultured in DMEM supplemented with 10% FBS and were supplemented with 100 U/mL penicillin and 100 μg/mL streptomycin and were cultured and humidified with 5% $CO_2$ at 37 °C. HPDEC were cultured in Keratinocyte growth medium.

**Confocal microscopy**

A confocal dish (35 mm dish with 20 mm bottom well, #1.5 glass) was used to prepare CLSM samples. For live-cell imaging, cells in exponential growth phase were seeded on the confocal dish at $1.0\times10^5$ cells per dish and incubated for 24h. After removing the culture medium, fresh medium containing the compound of interest was added to the cells for co-incubation at 37 °C in a humidified atmosphere of 5% $CO_2$ for the desired period. Afterwards, the nuclei of cells were stained with Hoechst 33342 for 10 minutes, and the samples were washed with 1 mL of Live Cell Imaging Solution four times to fully remove the residual Hoechst 33342.

For time-lapse live-cell imaging, cells in exponential growth phase were seeded on a confocal dish at $1.0\times10^5$ cells per dish and incubated for 24 h. The samples were washed with 1 mL of Live Cell Imaging Solution three times, and the nuclei were stained with Hoechst 33342 for 10 minutes. The samples were then washed with 1 mL of Live Cell Imaging Solution four times to remove the residual Hoechst 33342. The position of cells and the focal plane of the laser beam were determined using the fluorescence from the stained nuclei with a laser with 405 nm wavelength, and the Nikon Perfect Focus System was activated to prevent focus drift. The imaging solution in the confocal dish was replaced with fresh imaging solution containing the compound of interest. CLSM images of different channels were then recorded, with the timeseries interval set to be 30 seconds or no-delay. After a specified number of imaging cycles, the fluorescence images from both channels were saved for further analysis.

**Cell pretreated with inhibitors**

PANC-1 cells ($1.5\times10^5$ cells) were seeded in a confocal dish for 24 h to allow attachment. The culture media was then replaced with fresh medium containing inhibitor for palmitoylacyltransferases (2-BP, 20 μM) and the cells were incubated for 30 minutes at 37 ° C. Afterward, the cell medium was replaced with fresh medium containing CyMA-i for live-cell imaging.

**Immunocytochemistry**

The culture media was removed from cells, followed by three times washes with cold PBS. Cells were fixed with 4% paraformaldehyde for 10 minutes and permeabilized with 0.25% Triton X-100 in PBS for 10 minutes. Cells were then blocked with 3% BSA and 22.5 mg/mL of glycine in PBST for 1 hour at room temperature. Primary antibodies were diluted 1:200 in 1% BSA in PBST and incubated with the cells overnight at 4 °C. Cells were then incubated with Alexa Fluor 647-conjugated secondary antibodies (1:1000 dilution) for 1 hour at room temperature. Between each step, except after blocking, cells were washed three times with

PBS. The cells were stained with Hoechst 33342 for 10 minutes and washed three times with PBS or directly ready for CLSM imaging without staining.

**Palmitoylation of CyMA characterized by LC/HR-MS**

A total of $1.2\times10^7$ PANC-1 cells were treated with 1 μM of CyMA **1**, **2** or vehicle (DMSO) for 2 hours, and then washed with HEPES buffer twice. Cells were collected and centrifuged to obtain a pellet. The pellet was resuspended in 500 μL of HEPES buffer, followed by the addition of 1.5 mL of DCM and 1 mL of methanol. The mixture was allowed to sit for 10 minutes. Afterward, 0.5 mL DCM and 0.5 mL Tris-HCl (50 mM, pH 2.0) were added, and the tube was centrifuged to collect the organic phase. The organic phase was washed with buffer (1 mL methanol+1 mL Tris-HCl (50 mM, pH 2.0)) and centrifuged again to obtain the organic phase. The organic solvent was evaporated with $N_2$, and the remaining solid was dissolved in 150 μL of methanol and sent for LC/HR-MS analysis.

**Critical micelle concentration (CMC) measurement**

The critical micellar concentrations (CMC) were assessed utilizing the fluorescent spectra of pyrene motif on the peptides. Various concentrations of compounds were prepared in saturated pyrene in distilled water. By plotting the intensity ratios at 374 nm and 384 nm ($I_1/I_3$) for each concentration, the concentration at the intersection point of two fitted lines was identified as the CMC.

**Transmission electron microscopy (TEM) sample preparation**

Negative stain electron microscopy for most of the study due to its speed, convenience, and lower cost compared to cryo-EM. After placing 5 μL samples on 400 mesh copper grids coated with continuous thick carbon film (~35 nm) which was glowed discharged, we washed the grid with ddH2O and 2% UA (uranyl acetate). The sample loaded grid was stained with UA for 20 seconds. The residual UA was removed by filter paper and then dried in air. TEM images were obtained with FEI Morgagni 268 80 kV with a 1 k × 1 k AMT CCD camera.

**MTT assay**

The MTT assay was used to determine cell viability for cytotoxicity evaluation. Cells were seeded at $1\times10^4$ cells per well in 96-well plates for 24 h to allow attachment. Culture media were replaced with fresh culture media containing the compounds or compounds combined with different type cell death inhibitors at a series of concentrations. After 24, 48, and 72 hours, 10 μL of MTT solution (5 mg/mL) was added to each well, and the plate was incubated in the dark for 4 h at 37 °C. Then, 100 μL of 10% SDS-HCl was added to stop the reaction and dissolve the formazan. The absorbance at 595 nm was determined by a microplate reader. The assay was repeated three times, and the mean values of three measurements were plotted, with error bars representing standard deviation.

**Immunoblotting**

Cells were cultured to 80-90% confluency, lysed with 500 μL lysis buffer (containing protease inhibitor cocktail and phosphatase inhibitor cocktails) per 10 cm dish on ice, sonicated for 10 seconds, and subjected to three freeze-thawed cycles to collect cell lysates from various cell lines. The lysates were centrifuged at 12,000 rpm for 10 minutes at 4 °C, and the supernatant was collected. The proteins in the lysates were denatured by adding sample loading buffer and incubating at 95 °C for 10 minutes. The lysate samples were loaded onto precast gels for electrophoresis at 140 V for 40 minutes, followed by blotting onto PVDF membrane under 100 V for 90 minutes in ice bath. Membranes were blocked with blocking buffer for 1 hour at room temperature and incubated with primary antibody (1:1000 dilution) at 4 °C overnight. After washing with TBST three times (5 min per wash), the membrane was incubated with secondary antibody (1:10,000 dilution) for 1 hour at room temperature. After washing with TBST five times, chemiluminescent substrate was added and incubated for 1 minute. The membranes were then scanned using a blot scanner. Subsequently, the scanned membranes were stripped using Restor™ Western Blot Stripping Buffer, blocked, and incubated with the next primary antibody. Note: across replicate immunoblot experiments, total protein levels may show minor variation; however, after normalization, the ratios and the direction of change (increase or decrease) remain consistent.

**Cell spheroids generation**

PANC-1 Spheroids were generated using a low-adhesion U-bottom microplate, specifically the Thermo Scientific™ Nunclon™ Sphera™ 96U-well microplate (Cat. No. 174929). PANC-1 cells (10,000 cells per 50 µL) were seeded into each well and incubated in a cell incubator for 48 hours to generate spheroids.

**3D Cell Viability Assay**

The cytotoxicity of CyMA against the generated cell spheroids was tested using The CellTiter-Glo® 3D Cell Viability Assay, which measures ATP as an indicator of viability and generates a luminescent readout that is much more sensitive than colorimetric or fluorescence-based methods. Specifically, 100 µL of CellTiter-Glo® 3D Reagent was added into each well containing 100 μL of culture media with cell spheroids. The contents were mixed vigorously for 5 minutes to induce cell lysis, followed by a 25-minute incubation at room temperature to stabilize the luminescent signal. Luminescence was recorded using a microplate reader.

**Drug Resistance Test**

PANC-1 cells were divided into two culture dishes. Group 1: The PANC-1 cells were incubated with **1** (100 nM) for 24 hours, and the media was replaced with fresh media for 2 more days to allow the cells to expand. The cells were then subcultured and allowed to proliferate until they reached 80-90% confluency, followed by incubation with **1** (100 nM) to initiate the next cycle of cell stimulation. Group 2: The PANC-1 cells were incubated with vehicle (DMSO) instead of **1**, following the same procedure as a control. After 10 cell subculture cycles, we seeded stimulated cells in from Group 1 and unstimulated PANC-1 cells Group 2 in 96-well plates at $1\times10^5$ cells/well for 24 hours. The cells were then treated with **1** for 24 hours, and the cell viability was measured using the MTT assay.

**Quantification and Data Analysis**

All graphs were created using Origin 2024. Quantification of fluorescence intensity was conducted using Fiji. Error bars represent s.d. unless otherwise noted. For comparison between two groups, *p* values were determined using two-tailed Student's t-tests. The person's R value was calculated using Coloc 2 plugin in Fiji. All cell-based or cell-free experiments were repeated two to three times. Synergism effects are tested by Synergy Finder.[1]

### Supplementary Figures

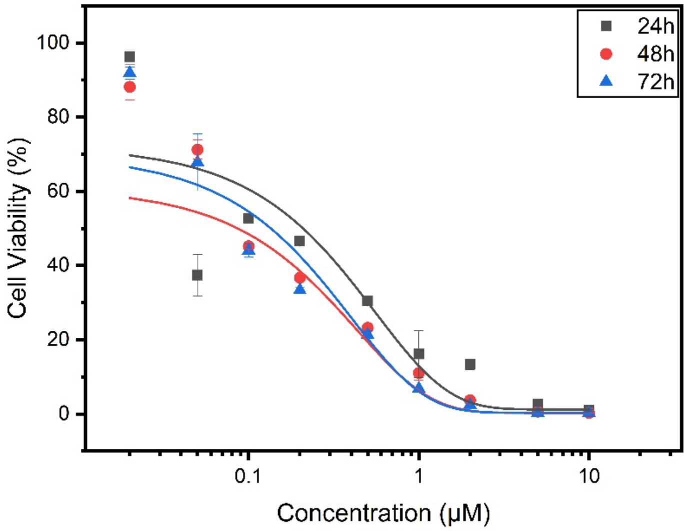


**Supplementary Figure S1.** The cell viability of PANC-1 cells treated by a serial concentration of Stb-$^{D}$Phe-$^{D}$Phe-SAc (**1**) for 24, 48 and 72 h.

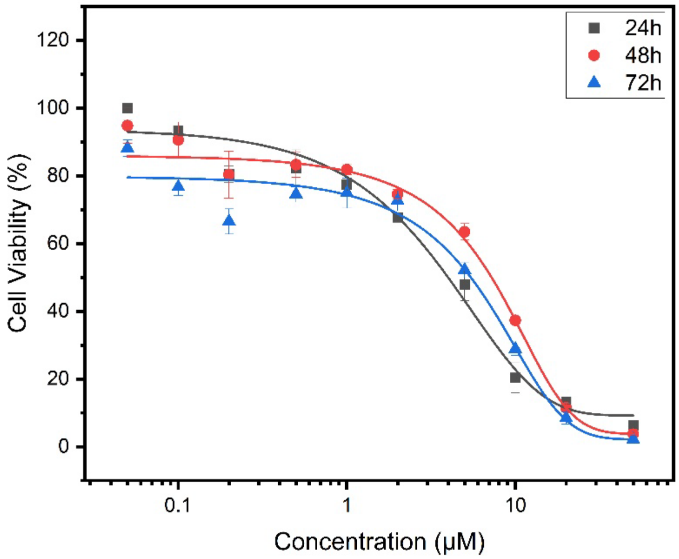


**Supplementary Figure S2.** The cell viability of PANC-1 cells treated by a serial concentration of Stb-$^{D}$Dap(NBD)-$^{D}$Phe-SAc (**2**) for 24, 48 and 72h.

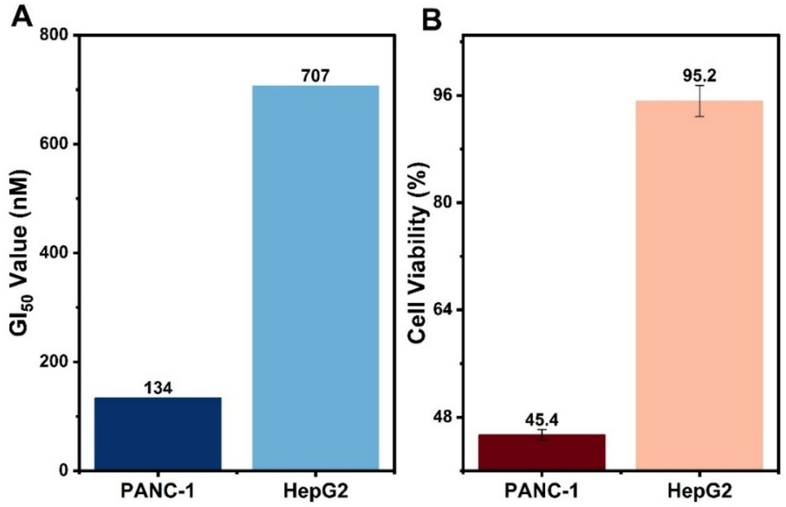


**Supplementary Figure S3.** (A) The GI$_{50}$ Values of Stb-$^{D}$Phe-$^{D}$Phe-SAc (**1**) towards PANC-1 and HepG2 cell lines at 24 h. (B) The cell viabilities of PANC-1 and HepG2 cells treated by 100 nM Stb-$^{D}$Phe-$^{D}$Phe-SAc (**1**) for 24h.

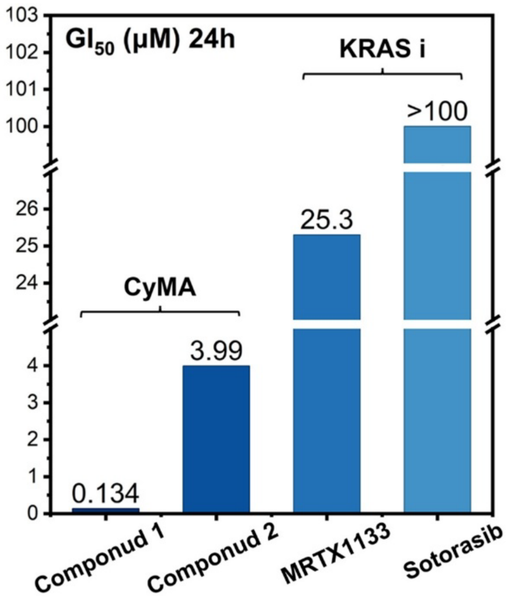


**Supplementary Figure S4.** GI$_{50}$ values of **1** and **2** compared with commercial KRAS inhibitors MRTX1133 (G12D mutation) and Sotorasib (G12C mutation) towards PANC-1 (24 h).

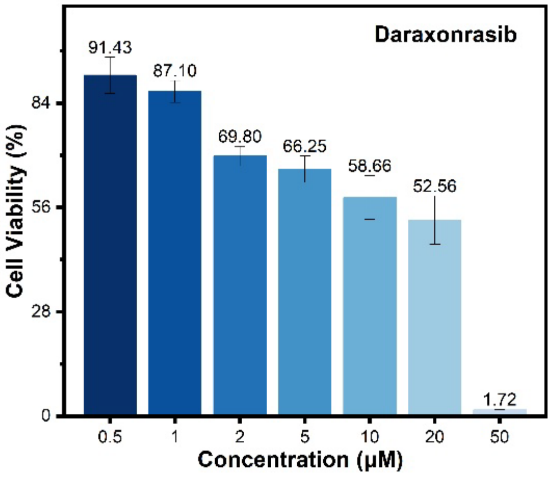


**Supplementary Figure S5.** The cell viability of PANC-1 cells treated by a serial concentration of Daraxonrasib for 24h.

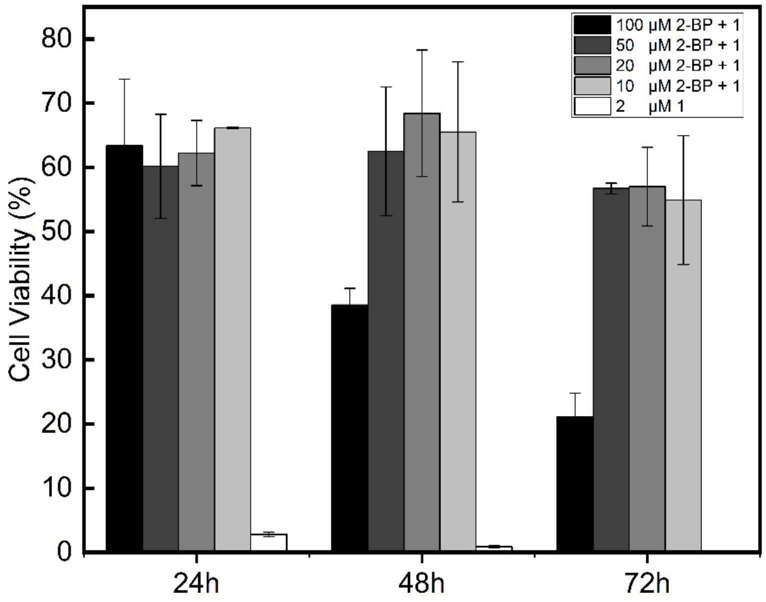


**Supplementary Figure S6.** The cell viability of PANC-1 cells treated by 2 μM Stb-$^{D}$Phe-$^{D}$Phe-SAc (**1**) combined with a serial concentration of 2-BP for 24, 48 and 72h.

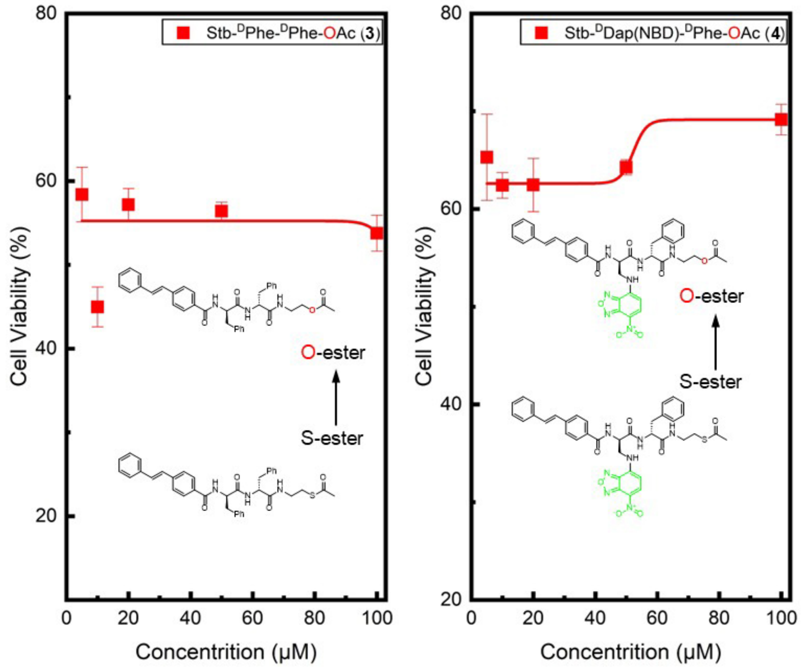


**Supplementary Figure S7.** The cell viability of PANC-1 cells treated by a serial concentration of Stb-DPhe-DPhe-OAc (**3**) and Stb-DDap(NBD)-DPhe-OAc (**4**) for 24, 48 and 72h.

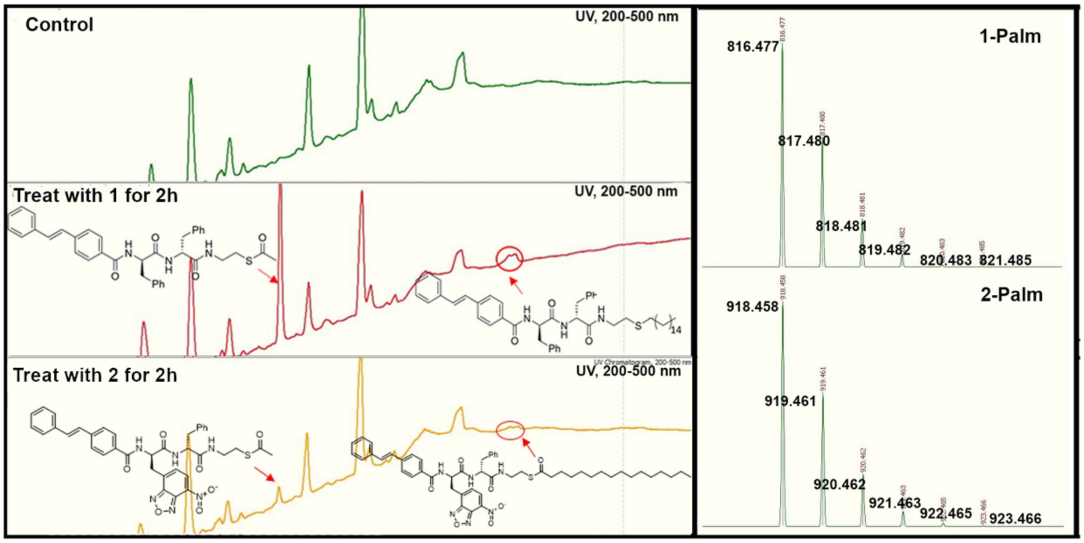


**Supplementary Figure S8.** High-resolution LC-Mass spectrum results. LC traces (left) of PANC-1 cell extracts treated w/o compound **1** or **2** for 2h. Isotope pattern (right) for compound **1-Palm** and **2-Palm**.

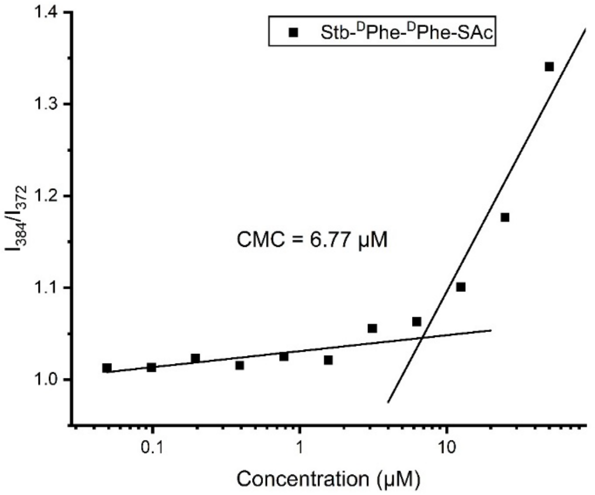


**Supplementary Figure S9.** Critical Micelle Concentrations of compound **1** dissolved in pyrene.

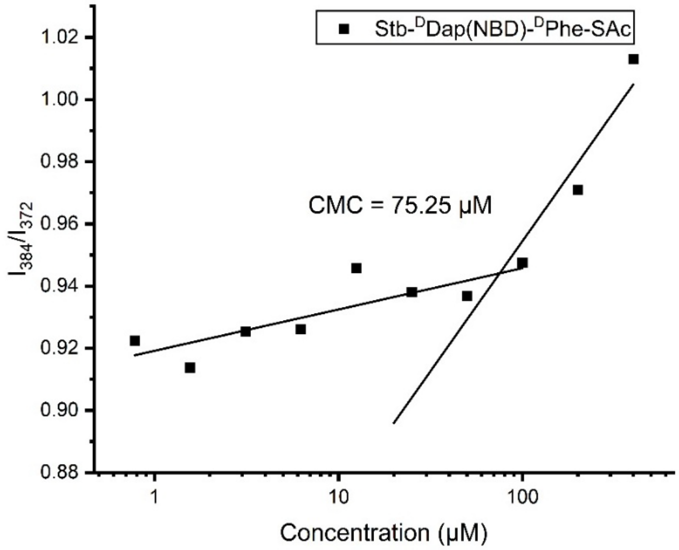


**Supplementary Figure S10.** Critical Micelle Concentrations of compound **2** dissolved in pyrene.

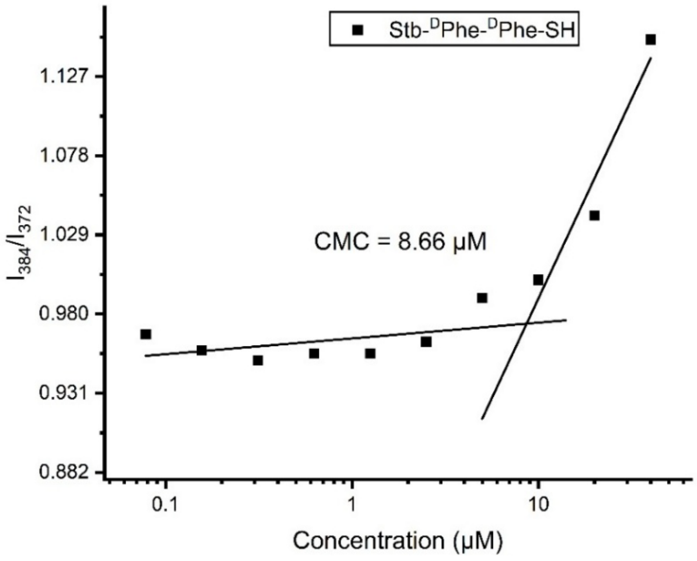


**Supplementary Figure S11.** Critical Micelle Concentrations of compound **1-SH** dissolved in pyrene.

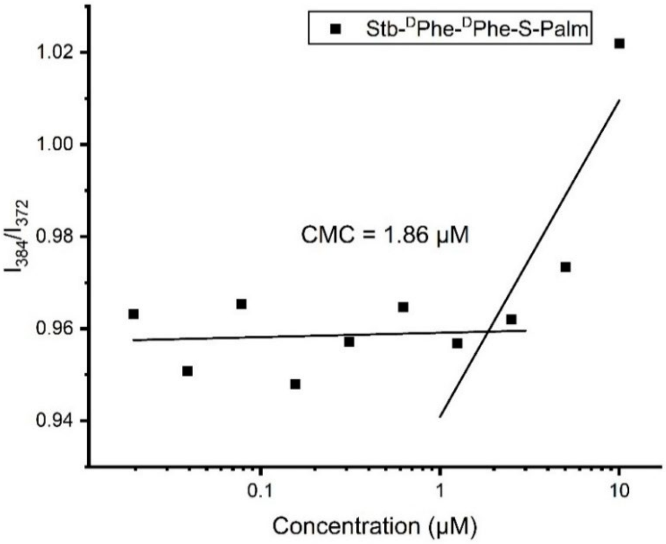


**Supplementary Figure S12.** Critical Micelle Concentrations of compound **1-Palm** dissolved in pyrene.

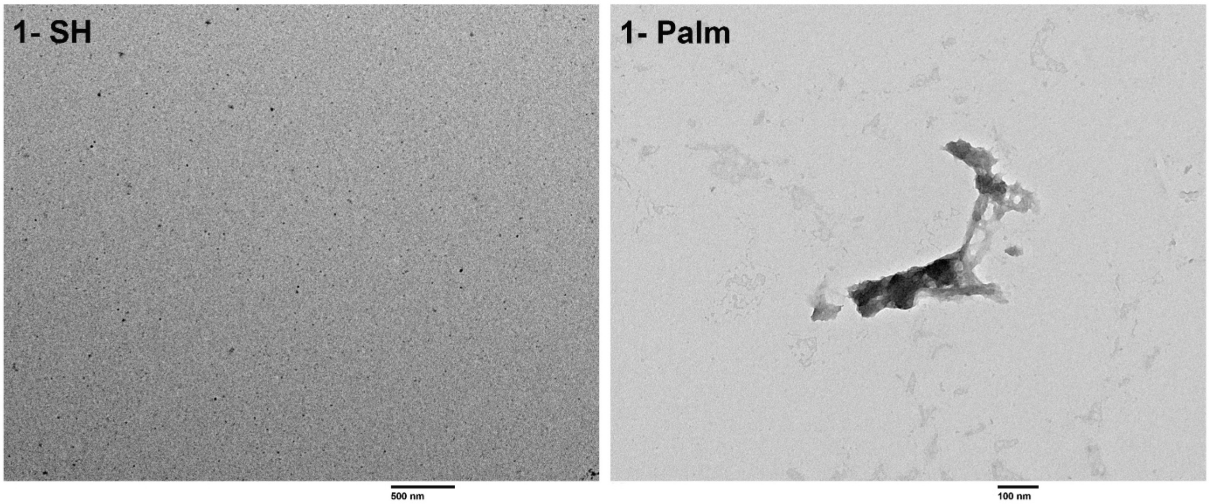


**Supplementary Figure S13.** TEM images of 2 μM **1-**SH (**5**) (left), 2 μM **1-**Palm (**6**) (right) in distilled water. Scale bars are 500 nm and 100 nm respectively.

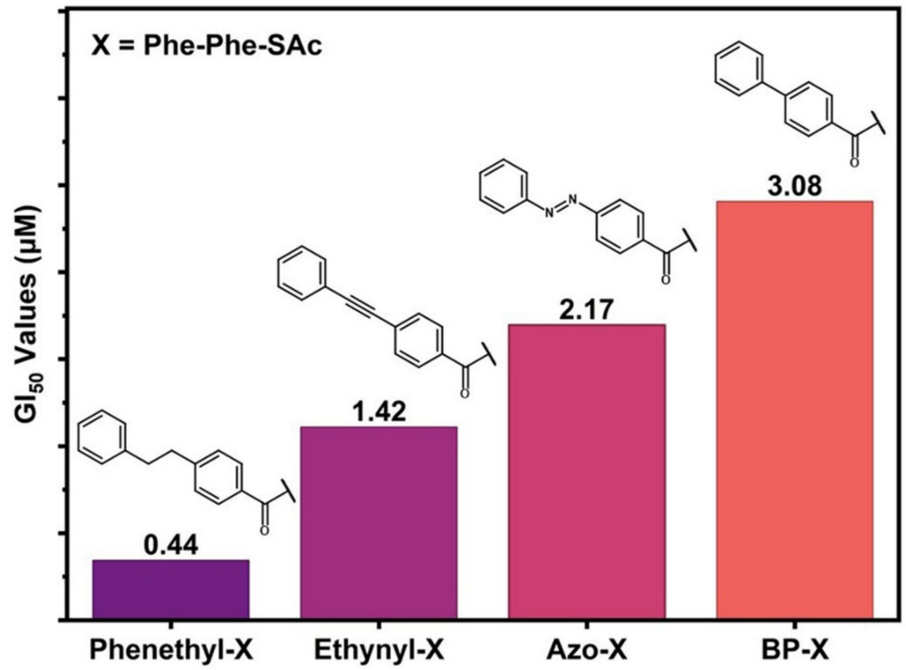


**Supplementary Figure S14.** $GI_{50}$ values of derivatives with different caping groups towards PANC-1 cell line for 24h.

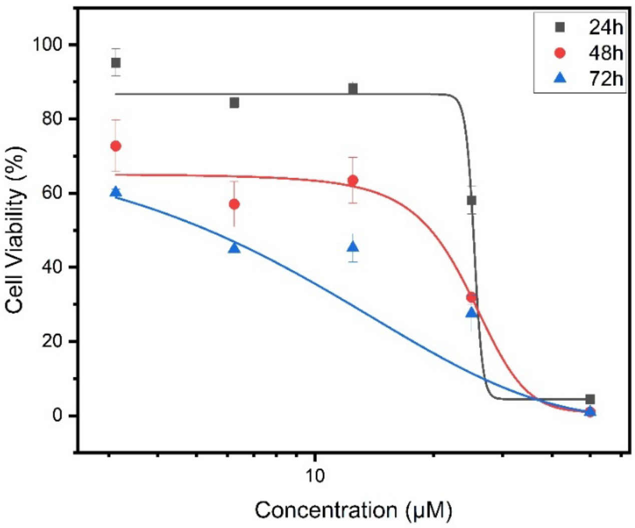


**Supplementary Figure S15.** The cell viability of PANC-1 cells treated by a serial concentration of MRTX1133 for 24h, 48h and 72h.

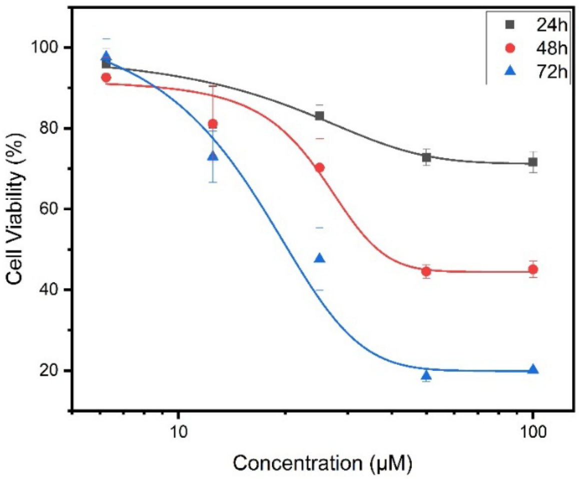


**Supplementary Figure S16.** The cell viability of PANC-1 cells treated by a serial concentration of Sotorasib for 24h, 48h and 72h.

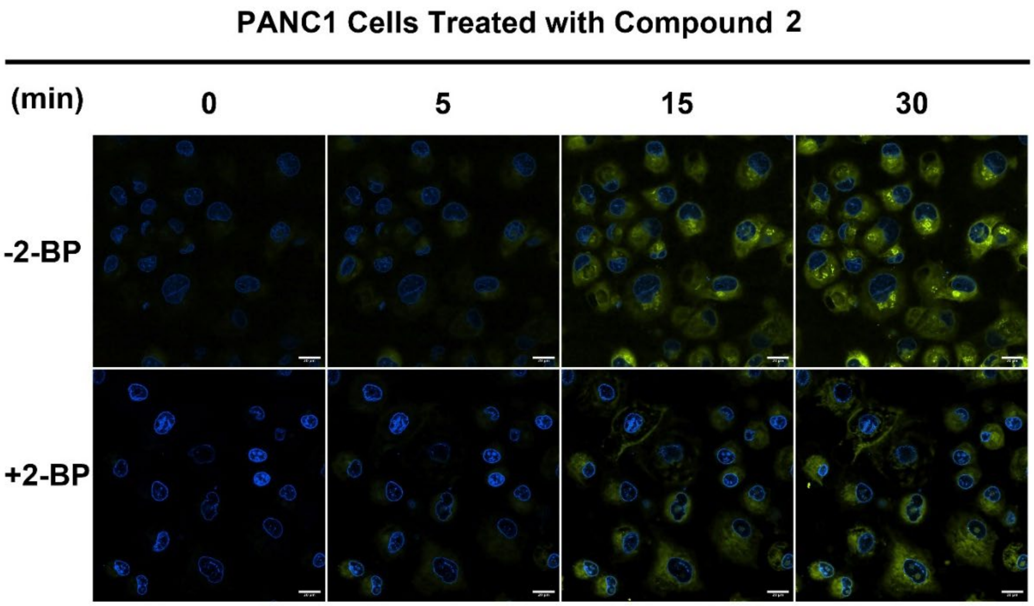


**Supplementary Figure S17.** CLSM images of PANC-1 cells treated with **2** (2 μM; 0, 5, 15, 30 min) with or without 2-bromohexadecanoic acid (2-BP) at concentration of 20 μM. Nuclei were stained by Hoechst 33342. Scale bar = 20 μm.

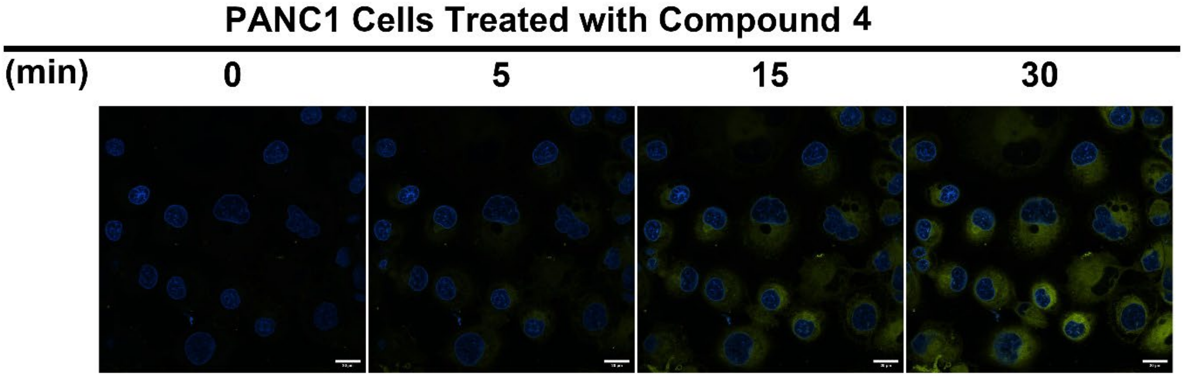


**Supplementary Figure S18.** CLSM images of PANC-1 cells treated with **4** (2 μM; 0, 5, 15, 30 min). Nuclei were stained by Hoechst 33342. Scale bar = 20 μm.

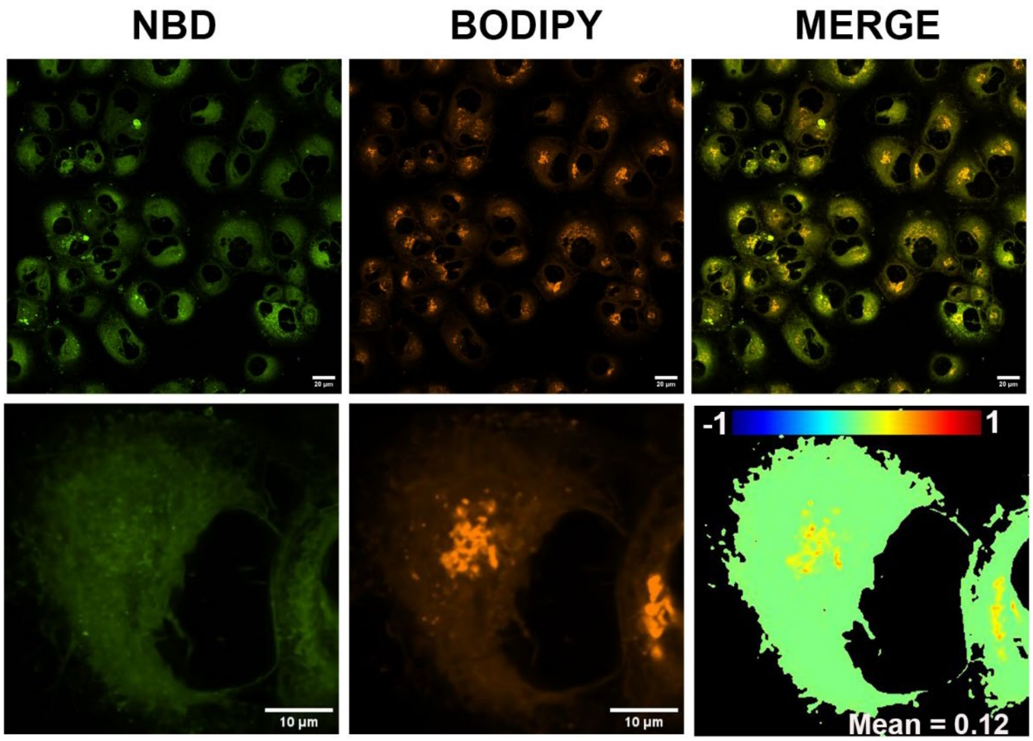


**Supplementary Figure S19.** CLSM images of PANC-1 cells treated with **4** (10 μM) combined with or without Golgi tracker for 2 h. Scale bar = 20 μm.

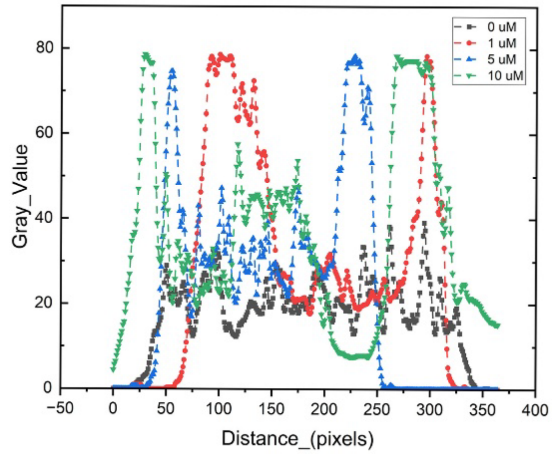


**Supplementary Figure S20.** Gray Value along the arrow of KRAS's ICC staining to the ratio of KRAS distributed in plasma membrane to cytosol changing after the treatment with or without **1** in Figure 2F.

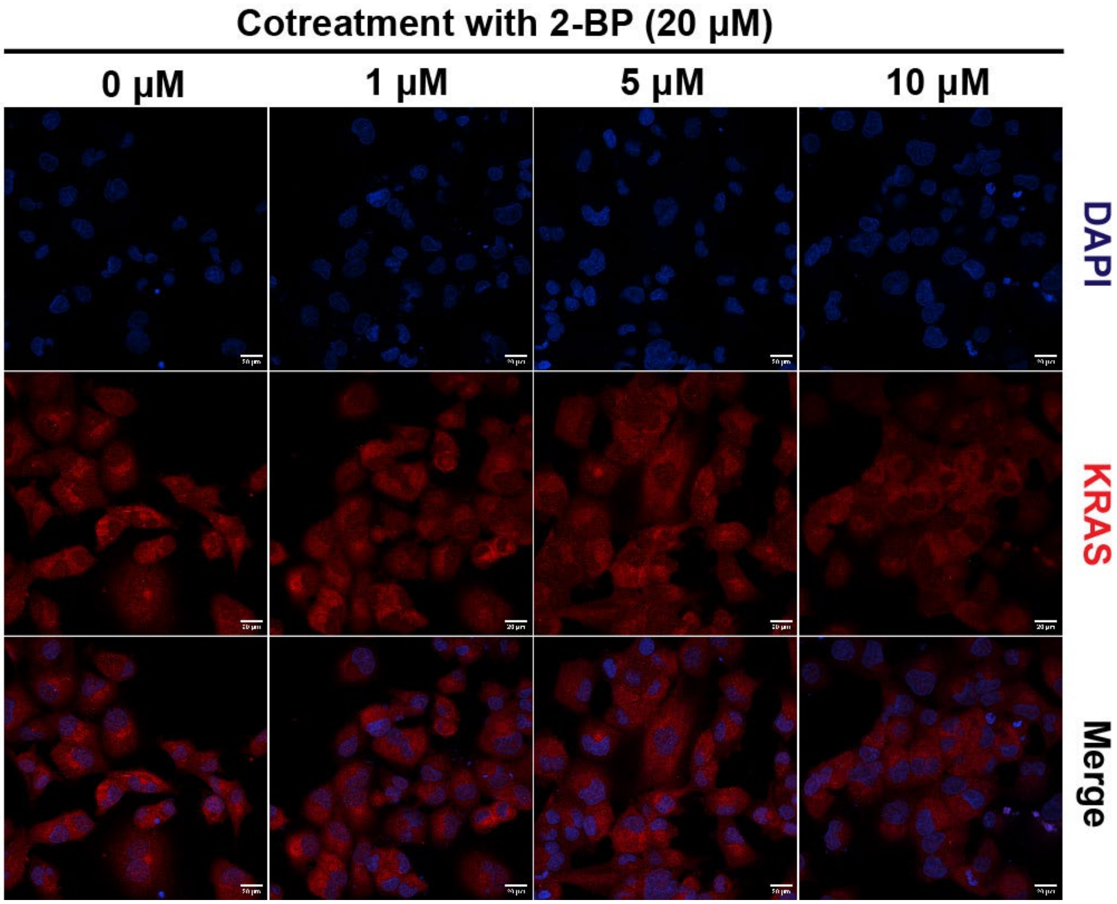


**Supplementary Figure S21.** Immunofluorescence images showing KRAS localization in PANC-1 cells after Cotreatment with 2-BP (20 μM) and **1** at the indicated concentrations for 12 h. KRAS was detected with an Alexa Fluor 647-conjugated secondary antibody, and nuclei were stained with Hoechst 33342. Scale bar = 20 μm.

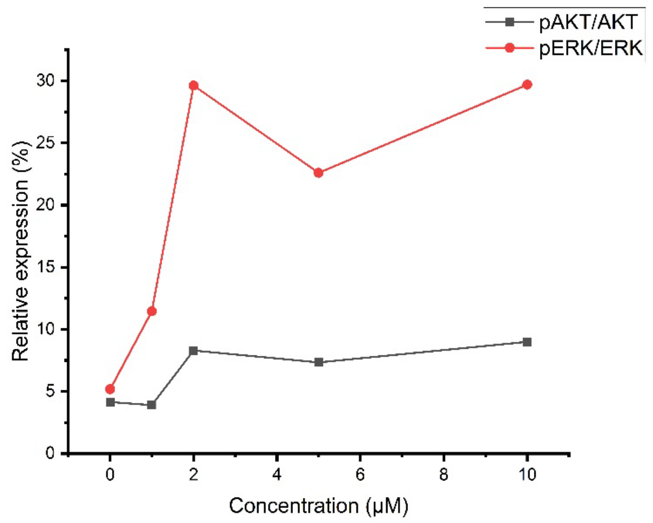


**Supplementary Figure S22.** The quantification of relative expression of pAKT/AKT and pERK/ERK in PANC-1 cells after being treated with different concentrations of **1** for 12h.

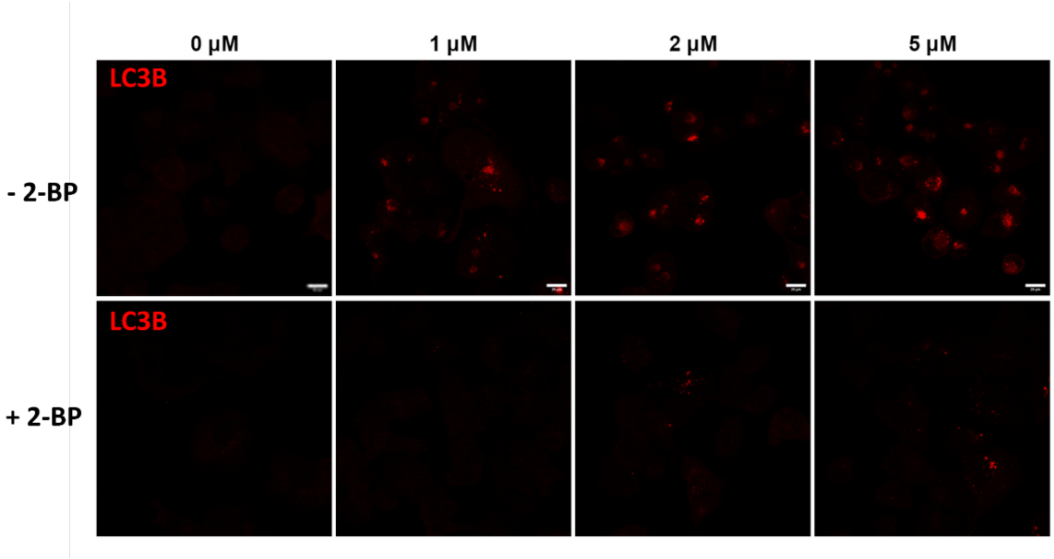


**Supplementary Figure S23.** Intracellular distributions of LC3B were determined by immunostaining with LC3B antibody of PANC-1 cells after pretreatment with or without compound **1** in 1, 2, 5 μM combined with or without 2-BP (20 μM) for 12h. Alexa Fluor 647 was used as second antibody (Red). Scale bar = 20 μm.

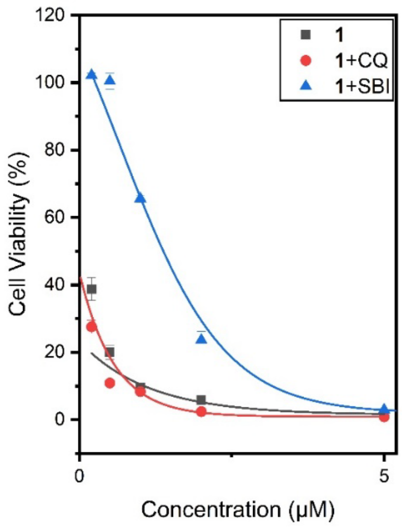


**Supplementary Figure S24.** The cell viability of PANC-1 cells treated by a serial concentration of Stb-DPhe-DPhe-SAc (**1**) combined with CQ (20 μM) or SBI-0206965 (5 μM) for 24h.

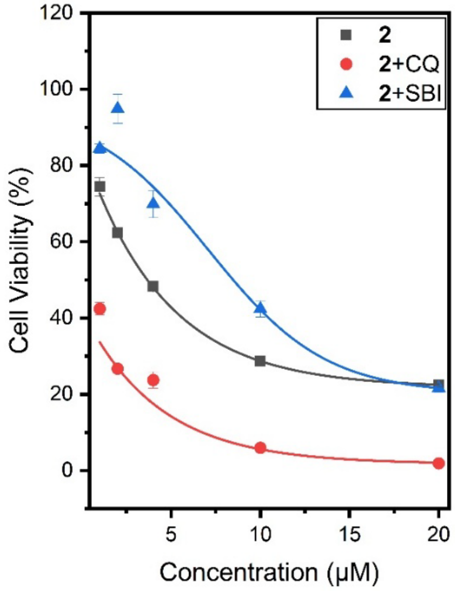


**Supplementary Figure S25.** The cell viability of PANC-1 cells treated by a serial concentration of Stb-DDap(NBD)-DPhe-SAc (**2**) combined with CQ (20 μM) or SBI-0206965 (5 μM) for 24h.

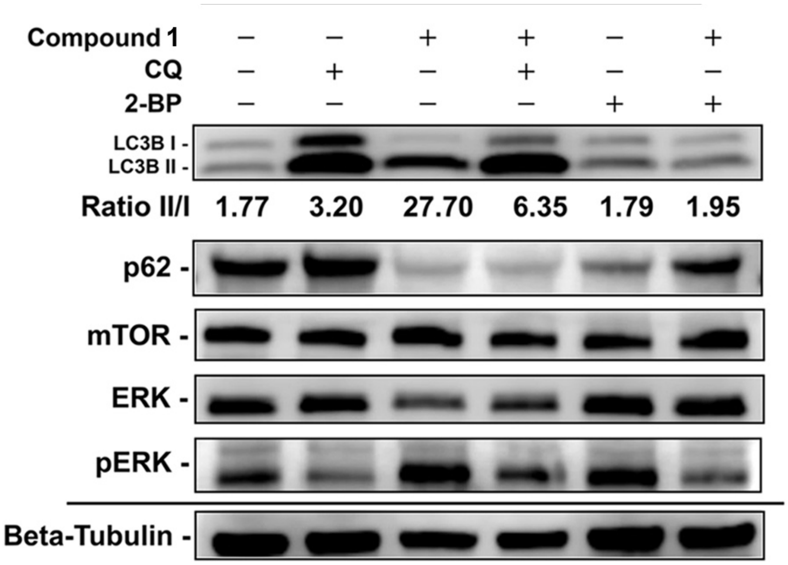


**Supplementary Figure S26.** Immunoblotting of Proteins after treatment with **1** (1 μM) alone or in combinations with CQ (20 μM) or 2-BP (20 μM) for 12 h.

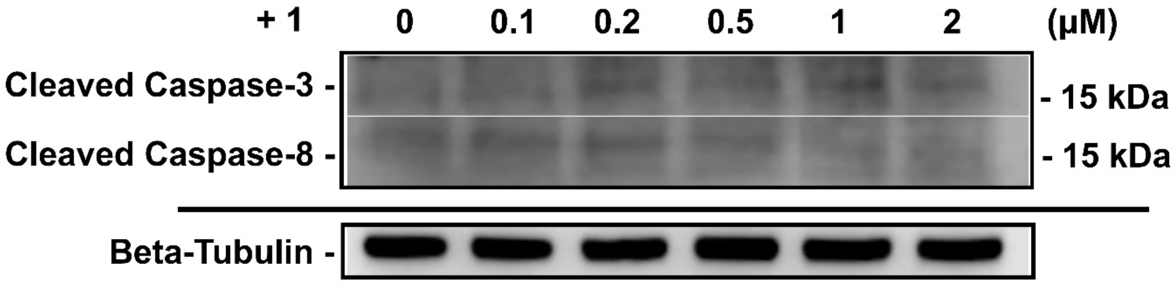


**Supplementary Figure S27.** Immunoblotting of cleaved-caspase-3 and cleaved-caspase-8 in PANC-1 cells treated with **1** at different concentrations for 12 h.

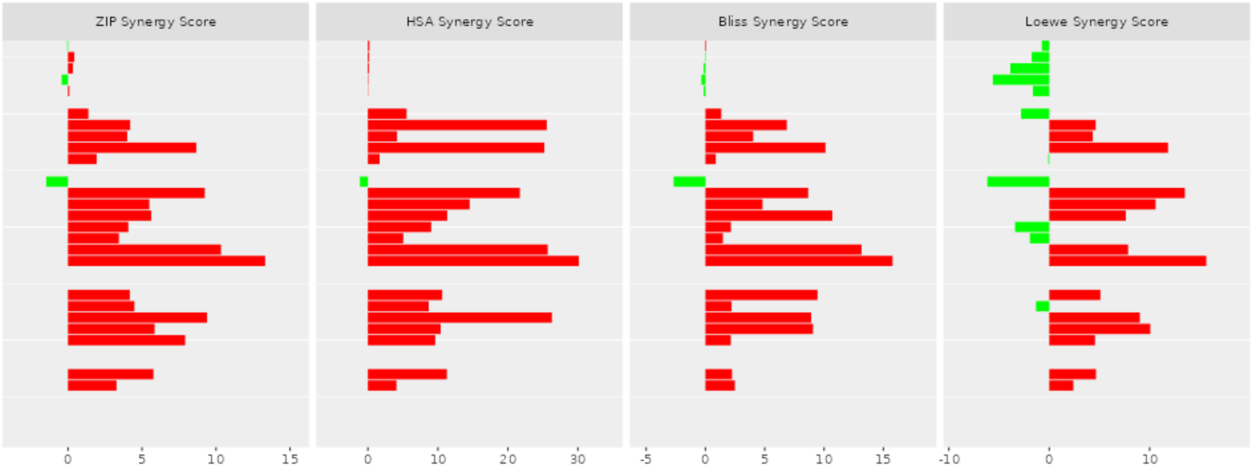


**Supplementary Figure S28.** Comparison of synergy scores obtained from different reference models. Synergy scores for all concentration combinations of 1 and BCI in PANC-1 cells were ranked and analyzed using the ZIP, HSA, Bliss, and Loewe models. The majority of combinations exhibited positive synergy scores, indicating cooperative anti-proliferative effects between the two agents.

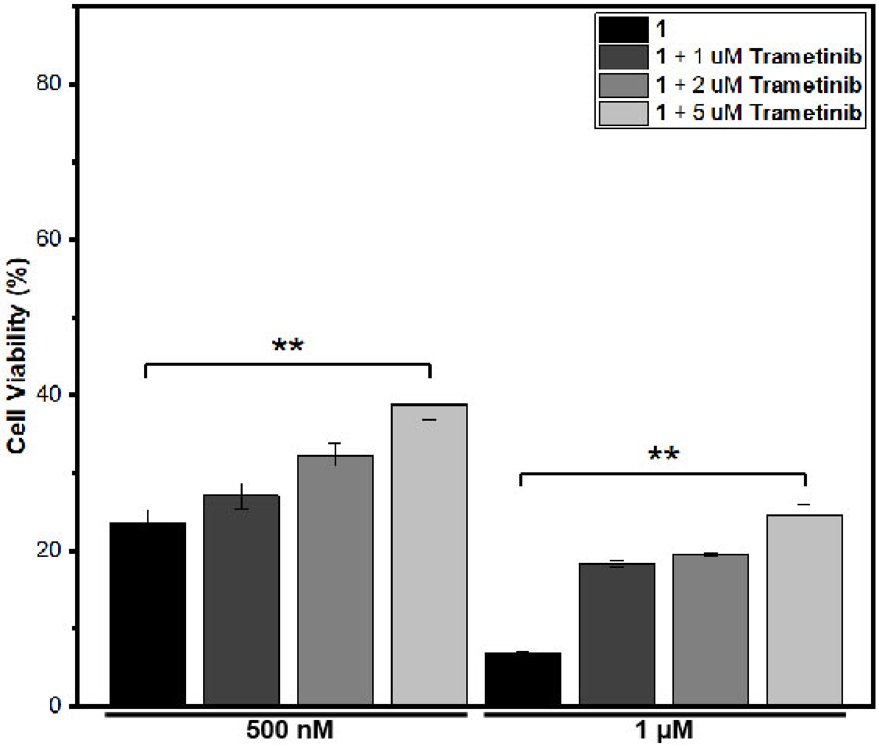

**Supplementary Figure S29.** The cell viability of PANC-1 cells treated by Stb-$^{D}$Phe-$^{D}$Phe-SAc (**1**) combined with a serial concentration of commercial MEK inhibitor trametinib for 24h. $^{**}P < 0.01$, n=3.

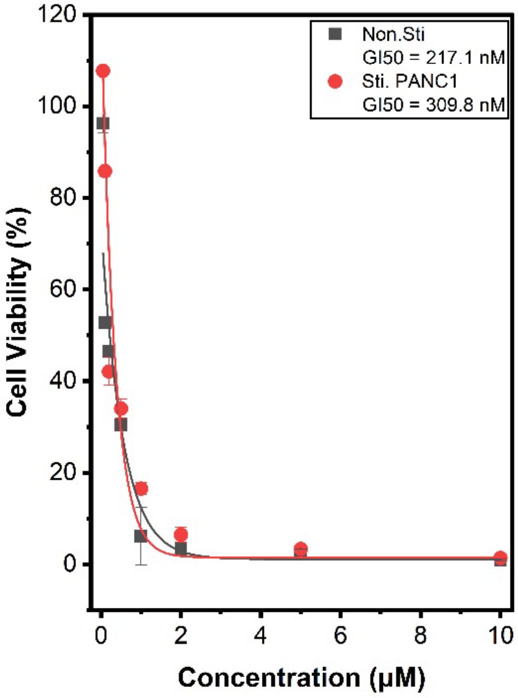


**Supplementary Figure S30.** The cell viability of stimulated/non-stimulated PANC-1 cells after 24h treatment of Stb-$^{D}$Phe-$^{D}$Phe-SAc (**1**) (n = 3).

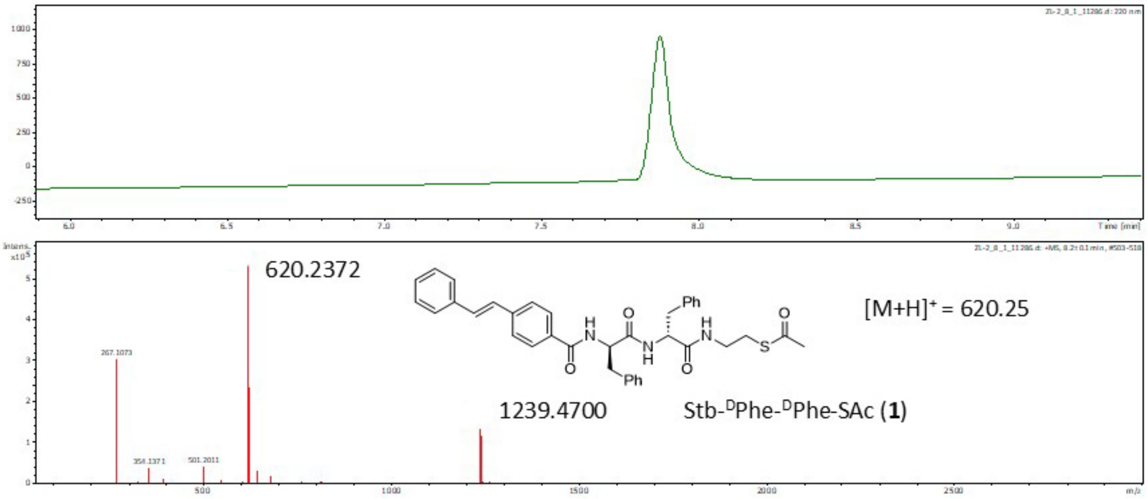


**Supplementary Figure S31.** The LC-HRMS of the synthesized compound **1**.

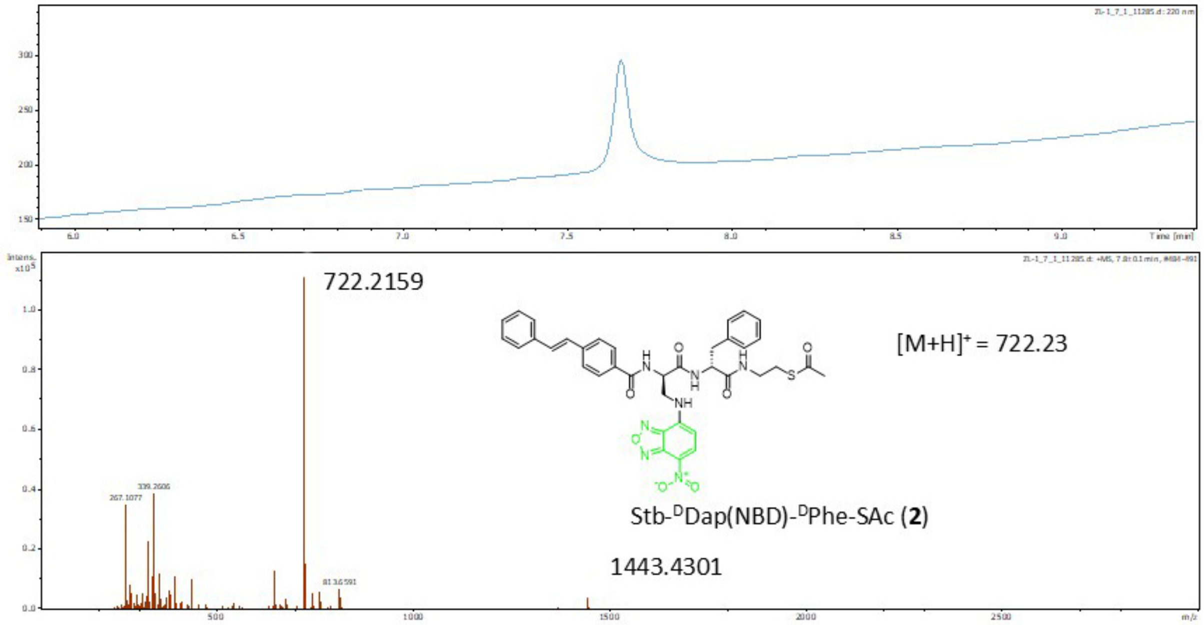


**Supplementary Figure S32.** The LC-HRMS of the synthesized compound **2**.

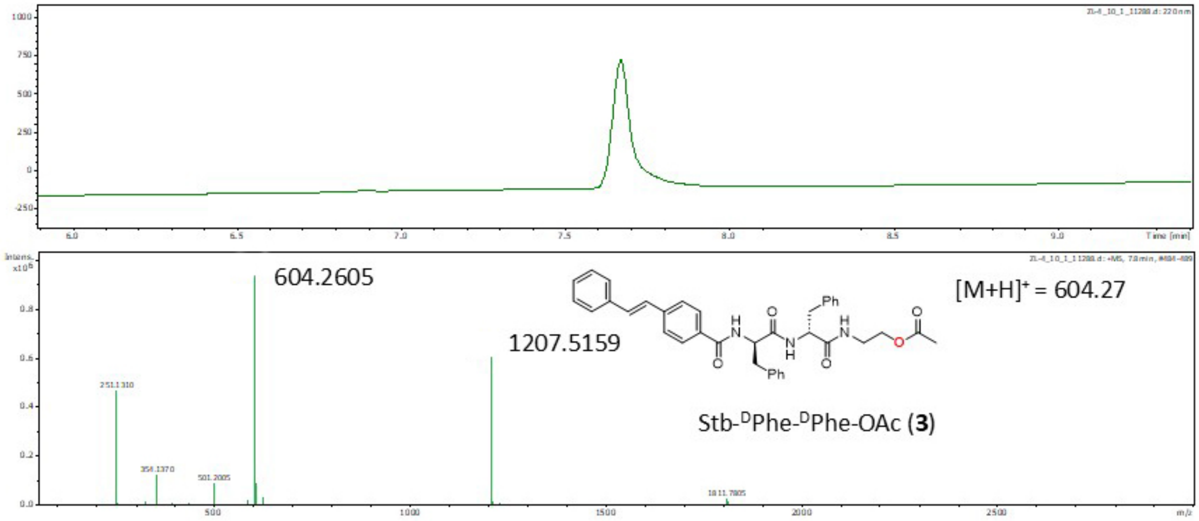


**Supplementary Figure S33.** The LC-HRMS of the synthesized compound **3**.

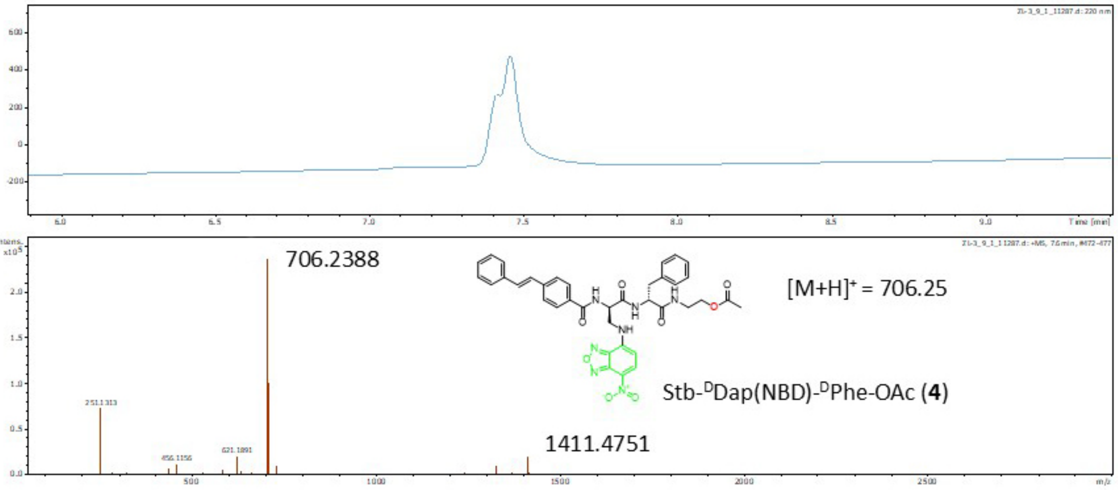


**Supplementary Figure S34.** The LC-HRMS of the synthesized compound **4**.

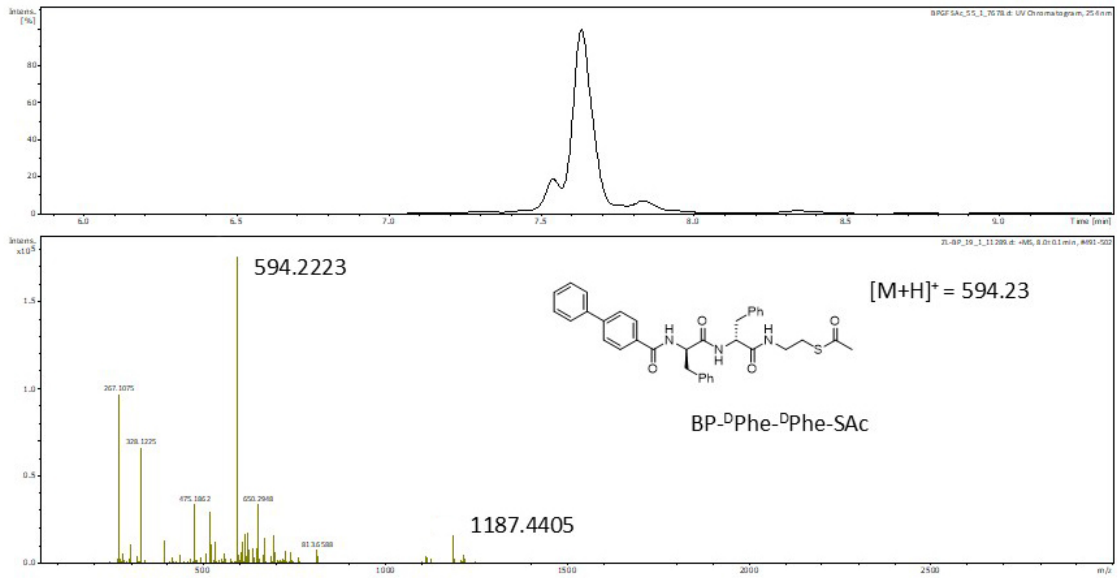


**Supplementary Figure S35.** The LC-HRMS of the synthesized compound BP-$^{D}$Phe-$^{D}$Phe-SAc.

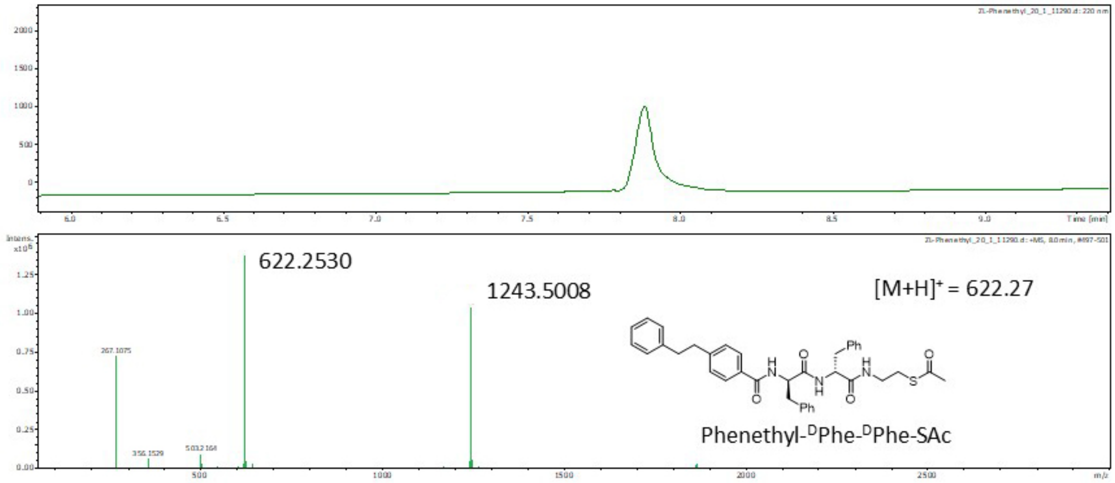


**Supplementary Figure S36.** The LC-HRMS of the synthesized compound Phenethyl-$^{D}$Phe-$^{D}$Phe-SAc.

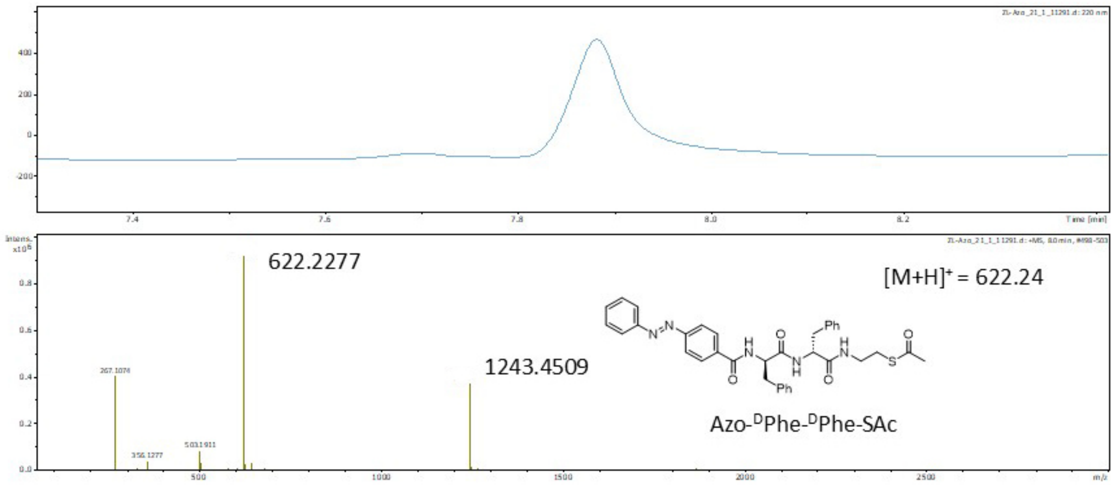


**Supplementary Figure S37.** The LC-HRMS of the synthesized compound Azo-DPhe-DPhe-SAc.

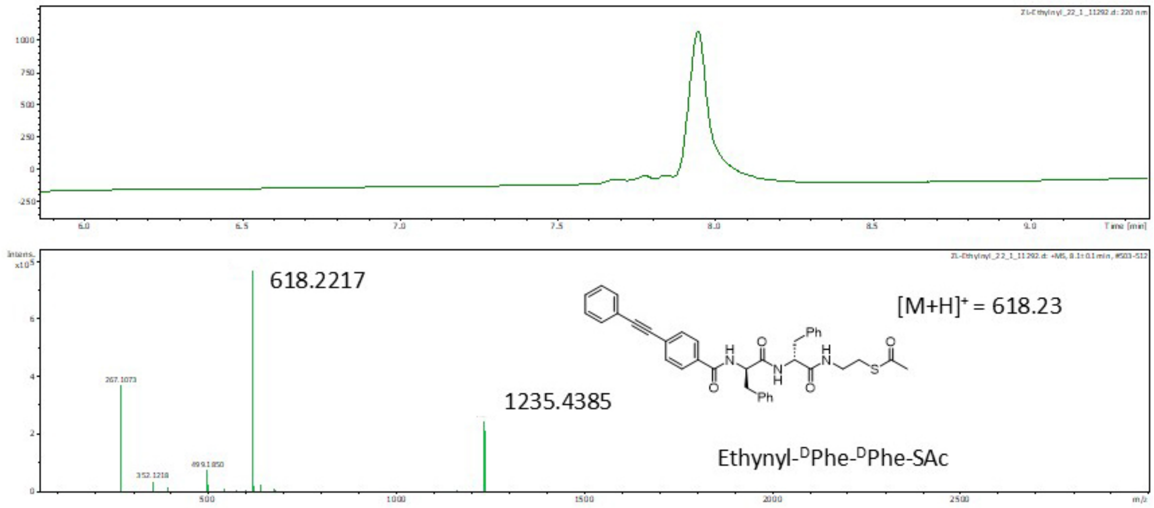


**Supplementary Figure S38.** The LC-HRMS of the synthesized compound Ethynyl-DPhe-DPhe-SAc.

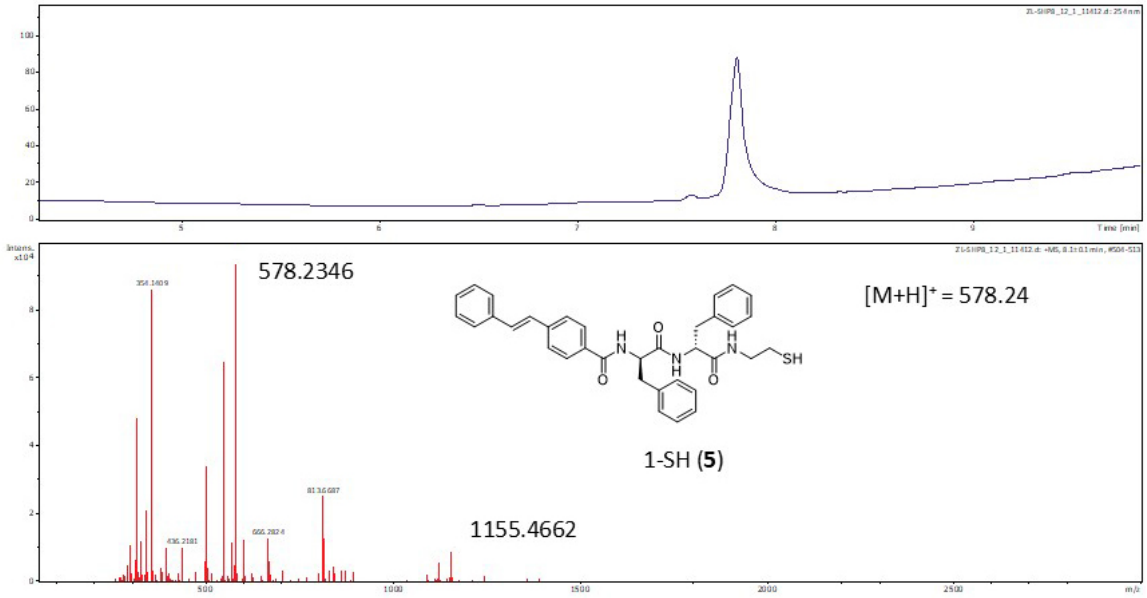


**Supplementary Figure S39.** The LC-HRMS of the synthesized compound **1-**SH.

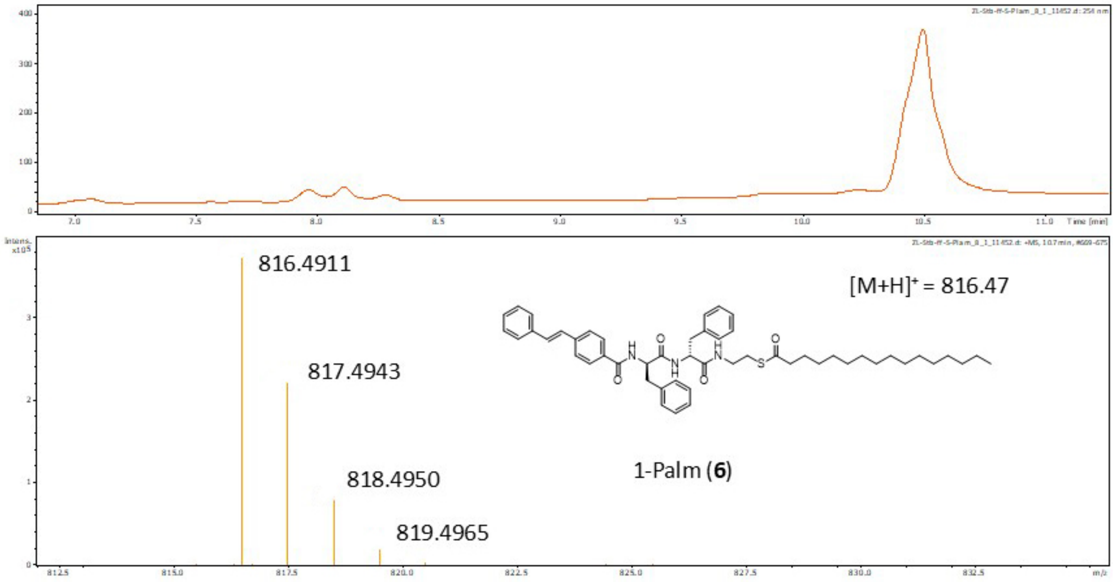


**Supplementary Figure S40.** The LC-HRMS of the synthesized compound **1-**Palm.
